# Nsg2 deficiency attenuates intestinal inflammation and colitis-associated tumorigenesis through microbiome-dependent metabolic remodeling

**DOI:** 10.64898/2026.11.23.767945

**Authors:** Gwanghee Kim, Kee Young Chung, Hosun Yu, Kee Jae Cho, Hyunsun Park, Jong Pil Lim, Byeong Gwan Kim, Joo Sung Kim, Seong-Joon Koh

**Affiliations:** Biomedical Research Institute, Seoul National University Hospital, Seoul, Republic of Korea; Department of Internal Medicine and Liver Research Institute, Seoul National University College of Medicine, Seoul, Republic of Korea; Laboratory of Intestinal Mucosa and Skin Immunology, Seoul National University College of Medicine, Seoul, Republic of Korea; Department of Biomedical Science, Seoul National University College of Medicine, Seoul, Republic of Korea; Department of Dermatology, Seoul National University College of Medicine, Seoul, Republic of Korea

**Keywords:** Nsg2, colitis, colorectal cancer, intestinal inflammation, microbiota

## Abstract

**Background & Aims:** Neuron-Specific Gene 2 (NSG2) is a neuronally expressed protein that facilitates excitatory synaptic transmission through the regulation of AMPA receptor trafficking. While its function has been characterized in the central nervous system, whether NSG2 plays a role in enteric neuronal signaling and intestinal pathology has yet to be explored. We therefore investigated whether Nsg2 deficiency modulates intestinal inflammation and tumorigenesis through gut microbiome-dependent mechanisms.

**Methods:** Dextran sulfate sodium (DSS)-induced colitis and azoxymethane/DSS-induced colitis-associated cancer (DSS-CAC) models were established in *Nsg2*^-/-^ and wild-type (WT) mice. Disease severity was evaluated using disease activity index (DAI), histopathological scoring, and tumor burden assessment. To investigate the potential contribution of the gut microbiota to genotype-associated phenotypes, WT and *Nsg2*^-/-^ mice were co-housed for six weeks prior to the experiment. Gut microbial composition and predicted functional pathways were analyzed using 16S rRNA gene sequencing (V3–V4 region) and PICRUSt2, respectively.

**Results:** Separately housed *Nsg2*^-/-^ mice exhibited significantly attenuated DSS-induced colitis, with reduced body weight loss, lower DAI scores, and preserved colonic architecture. In the DSS-CAC model, *Nsg2*^-/-^ mice developed significantly fewer and smaller tumors with suppressed Wnt/β-catenin signaling. Under co-housing conditions, no statistically significant differences in colitis severity or tumor burden were detected between genotypes, supporting a possible contribution of the gut microbiota to the phenotypes observed in separately housed mice. 16S rRNA sequencing identified distinct microbial communities in *Nsg2*^-/-^ mice, characterized by greater alpha diversity and enrichment of protective taxa, including Butyricicoccus and Lachnospiraceae. Functional profiling revealed upregulation of methane metabolism, porphyrin biosynthesis, and one-carbon metabolism pathways.

**Conclusions:** Nsg2 deficiency confers protection against intestinal inflammation and colitis-associated tumorigenesis, accompanied by alterations in gut microbial composition and suppression of Wnt/β-catenin signaling. These findings identify NSG2 as a novel modulator at the intersection of enteric neuronal signaling, gut microbiota, and mucosal immunity, positioning it as a potential therapeutic target for IBD and inflammation-driven colorectal cancer.

## Introduction

Inflammatory bowel disease (IBD), encompassing ulcerative colitis (UC) and Crohn’s disease, represents a chronic inflammatory disorder of the gastrointestinal tract with rising global incidence. The prevalence has markedly increased among young adults in Asia, particularly in Korea, Japan, and China, imposing substantial socioeconomic burdens.^1^ While the development of biological agents and small-molecule therapies has significantly improved patient outcomes, a considerable proportion of patients remain refractory to treatment or eventually lose therapeutic responsiveness.^2^ These therapeutic limitations underscore the urgent need for novel treatment strategies based on a deeper understanding of IBD pathophysiology.

Accumulating evidence has revealed a bidirectional relationship between IBD and neuropsychiatric disorders. Patients with IBD exhibit significantly higher risks of developing depression, anxiety, and neurodegenerative conditions such as dementia compared to the general population.^3,4^ These clinical observations support the emerging concept of the gut-brain axis, a complex bidirectional communication network linking the central nervous system (CNS), enteric nervous system (ENS), immune system, and gut microbiota.^5^

Within this axis, intestinal immune cells and gut microorganisms engage in dynamic crosstalk with both the CNS and ENS.^6^ The vagus nerve serves as a critical component of this network, forming synaptic connections with the ENS and modulating immune responses in the lamina propria through neurotransmitter signaling.^7^ Conversely, immune cells activated by microbial stimuli release pro-inflammatory cytokines, including TNF-α, IL-1β, and IL-6, which can traverse the blood-brain barrier and influence neuronal function by acting on receptors expressed in neurons and microglia.^6^ These interconnected pathways suggest that a systems-level understanding of gut-brain communication may be crucial for elucidating IBD pathogenesis.

Neuron-Specific Gene 2 (NSG2) has recently emerged as a key regulator of excitatory neurotransmission through its role in AMPA (α-amino-3-hydroxy-5-methyl-4-isoxazolepropionic acid) receptor trafficking and membrane expression during synaptic development.^8^ In the CNS, NSG2 is essential for maintaining excitatory synaptic signaling and neuronal plasticity. Loss-of-function mutations or genetic knockout of NSG2 result in impaired AMPA receptor trafficking and attenuated excitatory signaling, while NSG2 overexpression enhances AMPA receptor surface expression and synaptic excitability.^8^ Notably, increased AMPA receptor expression has been observed in the hippocampus of mice with TNBS-induced colitis, suggesting a potential connection between intestinal inflammation and altered excitatory neurotransmission.^9^ However, the specific contribution of NSG2 to intestinal inflammation and its interactions with the ENS and gut microbiota remain unexplored.

Here, we investigated the functional role of NSG2 in intestinal inflammation using *Nsg2* knockout mice. Our study reveals that NSG2 deficiency confers protection against both acute colitis and colitis-associated colorectal cancer through microbiome-dependent mechanisms. We demonstrate that *Nsg2* deletion alters gut microbial composition and enhances beneficial metabolic pathways, identifying NSG2 as a novel therapeutic target within the microbiota-gut-ENS axis for IBD treatment.

## Materials and Methods

### Animals

All mice used in this study were 7 to 8 weeks old, male C57BL/6 background mice and were maintained under SPF conditions in an animal facility at the Laboratory of Experimental Animal Research at Seoul National University Boramae Medical Center. Age-matched (7-8 weeks old) male wild-type (WT) C57BL/6 mice were purchased from Orient Bio Inc. (Seongnam, Republic of Korea) and served as controls. Mice were separated or cohoused for at least six weeks after weaning. All animal studies were approved by the Institutional Animal Care and Use Committee (IACUC) of Seoul National University Boramae Medical Center (IACUC approval number: 2021-0016). Animal care and experimental procedures were conducted in accordance with the National Institutes of Health Guide for Care and Use of Laboratory Animals and the Korean Animal Protection Act.

### 3D Cleared Tissue Imaging Analysis

Colonic tissue was harvested from anesthetized mice and immediately fixed in 4% paraformaldehyde (PFA). Tissue clearing was performed using a combined active and passive clearing approach. Briefly, fixed tissues were incubated in a decolorizing solution consisting of 5% CHAPS and 12.5% N-methyl diethanolamine in PBS for 72 hours, followed by active electrophoretic clearing using the X-Clarity™ tissue clearing system (Logos Biosystems, Cat. No. C30001) for 12 hours. Passive clearing was subsequently performed by incubation in SDS buffer (Logos Biosystems, Cat. No. C13001) at 58°C for 8 hours. Tissues were then washed in PBS (3 × 10 minutes) prior to immunolabeling.

For immunofluorescence staining, cleared tissues were incubated with primary antibodies against Tuj1 (1:200; Abcam, Cat. No. ab18207) and F4/80 (1:100; Invitrogen, Cat. No. MA1-91124) at 37°C for 72 hours. Following three washes in PBS for 10 minutes each, tissues were incubated with secondary antibodies donkey anti-rabbit IgG H&L Alexa Fluor 488 (1:500; Jackson ImmunoResearch, Cat. No. 711-545-152) and donkey anti-mouse IgG H&L Alexa Fluor 594 (1:500; Jackson ImmunoResearch, Cat. No. 715-585-150) at 37°C for 72 hours. After a final PBS wash (3 × 10 minutes), tissues were incubated in refractive index matching solution (Crayon Technologies, Cat. No. 50-3010) at 37°C for 48 hours.

Optical imaging was performed using an upright confocal microscope (C2+; Nikon Instruments). Three-dimensional image reconstruction and quantitative analysis were carried out using IMARIS software (v10.0.3; Oxford Instruments).

### Intestinal Permeability Assay

Intestinal permeability was assessed using fluorescein isothiocyanate (FITC)-dextran. Mice were fasted for 4–6 hours before oral administration of FITC-dextran (4 kDa; Sigma-Aldrich), with free access to water throughout the fasting period. Mice were housed in clean cages without bedding or food during fasting. FITC-dextran was dissolved in sterile 1× phosphate-buffered saline (PBS) at a concentration of 80 mg/mL, and mice were administered 200 μL of the solution by oral gavage. Four hours after administration, mice were euthanized and whole blood was collected. Serum was isolated by centrifugation at 5000 rpm for 10 minutes at 4°C and transferred to clean microcentrifuge tubes. Serum samples were loaded into black, opaque-bottom 96-well plates, with PBS included as a blank control. FITC-dextran fluorescence was measured using a microplate reader at excitation and emission wavelengths of 485 and 530 nm, respectively.

### Isolation of Lamina Propria Mononuclear Cells (LPMCs)

Lamina propria mononuclear cells (LPMCs) were isolated from colonic tissues using a modified protocol based on previously described methods. Briefly, colons were longitudinally opened and washed thoroughly with Hank’s Balanced Salt Solution (HBSS) supplemented with 1% fetal bovine serum (FBS). Tissues were cut into ∼0.5 cm² segments and incubated at 37°C for 20 minutes in HBSS containing 3 mM EDTA and 1 mM dithiothreitol (DTT) to remove the epithelial layer. The supernatant containing detached epithelial cells was filtered through a 100-μm cell strainer and discarded. Residual tissue fragments were finely minced and digested in α-minimum essential medium (MEM-α) supplemented with 10% FBS, 100 U/ml penicillin, 100 μg/ml streptomycin, 1 mg/ml collagenase IV (Biosharp, China), 1 mg/ml dispase II (Roche, Switzerland), and 0.5 mg/ml deoxyribonuclease I (Roche, Switzerland). Digestion was carried out for 60 minutes at 37°C in a shaking incubator. The resulting cell suspension was filtered through a 70-μm cell strainer (BD Biosciences, USA) and centrifuged at 300 × g for 10 minutes to collect LPMCs.

### Flow Cytometric Analysis of Immune Cell Populations

LPMCs were stained with fluorochrome-conjugated antibodies targeting surface markers to identify immune cell subsets, including T cells, B cells, neutrophils, and macrophages. The gating strategy is illustrated in Supplementary Figure S1. The following antibodies were used: Live/Dead cell (DAPI, Zomebie Green^TM^ Fixable Viabiltiy Kit), CD45 (BUV395, clone 30-F11), CD3 (APC-Cy7, clone 17A2), CD4 (BV650, clone GK1.5), CD8α (BV711, clone 53-6.7), Ly6G (PE, clone 1A8), Ly6C (PE-Cy7, clone AL-21), CD11b(BV421, clone M1/70), CD64 (APC, clone X54-5/7.1), MHC class II (BV786, clone M5/114.15.2). Surface staining was performed in PBS at 4°C for 30 minutes, followed by two washes with PBS. All antibodies were purchased from Abcam (UK) or BioLegend (USA), and staining protocols followed the manufacturers’ instructions. Flow cytometry was performed using a FACSAria flow cytometer (BD Biosciences, USA) to analyze immune cell populations within the lamina propria. Flow cytometry data were analyzed using FlowJo software (Tree Star, USA).

### DSS (Dextran Sodium Sulfate)-induced colitis

To induce colitis, mice were administered 3% (w/v) dextran sulfate sodium (DSS; colitis grade, M.W. 36–50 kDa; MP Biomedicals, Cat# 160110) in drinking water ad libitum for 6 or 7 days, followed by a 1- or 2-days recovery period with regular drinking water. Colitis severity was assessed using the Disease Activity Index (DAI), which ranges from 0 to 13 and is calculated based on body weight loss (<1%, 0; 1–5%, 1; 5–10%, 2; 10–15%, 3; 15–20%, 4; >20%, 5), stool consistency (normal, 0; loose stool, 2; diarrhea, 4), and rectal bleeding (no bleeding, 0; hemoccult-positive, 2; gross bleeding or perianal blood, 4). On day 6 or 7, mice were euthanized, and colon lengths were measured.

### Azoxymethane (AOM)/DSS-induced colitis-associated colon cancer

The colitis-associated colorectal cancer (CAC) model was established using azoxymethane (AOM) and dextran sulfate sodium (DSS), as previously described.^10^ Mice were intraperitoneally injected with AOM (12.5 mg/kg; Sigma–Aldrich, St. Louis, MO, USA) on day 0. Beginning on day 7, 3% (w/v) DSS was administered in drinking water for 5 consecutive days, followed by a 16-day recovery period with regular drinking water. This AOM/DSS cycle was repeated a total of three times. Body weight was recorded weekly in a blinded manner throughout the experimental period. On day 67 (Figure 3A) or 86 (Figure 5A), following completion of the third DSS cycle, mice were euthanized under isoflurane anesthesia. Colon tissues were harvested for macroscopic tumor assessment and histopathological analysis. This model enabled the evaluation of inflammation-driven tumorigenesis in the colon.

### Histological scoring

Colonic tissues were fixed in 10% neutral-buffered formalin, embedded in paraffin, sectioned, and stained with hematoxylin and eosin (H&E) for histological analysis. Colitis severity was evaluated using a standardized scoring system based on four criteria^11^: (1) inflammation severity (0: none; 1: mild; 2: moderate; 3: severe), (2) extent of inflammation (0: none; 1: mucosa; 2: mucosa and submucosa; 3: transmural), (3) crypt damage (0: none; 1: damage to the basal one-third of crypts; 2: damage to the basal two-thirds; 3: loss of entire crypts with intact surface epithelium; 4: complete loss of crypts and surface epithelium), and (4) percentage of tissue involvement (0: 0%; 1: 1–25%; 2: 26–50%; 3: 51–75%; 4: 76–100%). The total histological score was calculated by summing the individual parameter scores multiplied by the extent of tissue involvement.

### Immunohistochemistry (IHC)

Immunohistochemistry was performed as previously described¹² using primary antibodies against phosphorylated IκB kinase α/β (pIKK-α/β; Sigma–Aldrich, USA), β-catenin (Sigma– Aldrich, USA), and NSG2 (Abcam, UK). NSG2 immunostaining was performed on rectal tissue specimens from one patient with UC and one normal control (IRB approval no. 1104-066-358). Quantitative assessment of immunoreactivity followed a semi-quantitative visual scoring system, as described previously^11^. Staining intensity was evaluated based on the percentage of positively stained cells and graded as follows: 0 (no staining), 1+ (<10%), 2+ (10–30%), 3+ (31–60%), and 4+ (61–100%). For each marker, immunohistochemical intensity was determined in three randomly selected, nonadjacent tumor regions at ×400 magnification. To assess β-catenin nuclear translocation, the number of β-catenin–positive nuclei were counted in three high-power fields (HPFs) per slide.

### Stool DNA Extraction

Fecal samples were collected using sterile, individually packaged stool-sample containers (Mediclin®, Republic of Korea), and immediately frozen at −20 °C. Samples stored for longer than two weeks were transferred to −80 °C until further processing. Microbial genomic DNA was extracted using the E.Z.N.A.® Stool DNA Extraction Kit (Omega Bio-tek, USA) according to the manufacturer’s protocol. DNA was eluted in 50 μL of elution buffer, and its concentration and purity were assessed using a NanoDrop One UV–Vis spectrophotometer (Thermo Fisher Scientific, USA).

### 16S rRNA Gene Sequencing and Bioinformatic Analysis

The V3–V4 hypervariable regions of the bacterial 16S rRNA gene were amplified using the 341F (5′-CCTACGGGNGGCWGCAG-3′) and 805R (5′-GACTACHVGGGTATCTAATCC-3′) primer set (Cosmogenetech, Daejeon, Republic of Korea), in accordance with the Illumina 16S metagenomic library preparation protocol. Amplification was performed using OneTaq Hot Start 2X Master Mix (New England Biolabs, USA), and amplicons were quantified using the GenNext NGS Library Quantification Kit (Toyobo, Japan).

Sequencing was performed on an Illumina MiSeq platform using a 250 bp paired-end protocol (Illumina, USA). Raw reads were processed using QIIME2 (version 2024.10). Paired- end reads were trimmed, joined, and denoised with DADA2 to generate amplicon sequence variants (ASVs). Taxonomic classification was performed using the SILVA 132 reference database via the QIIME2-integrated classifier. Alpha diversity metrics (Chao1, Simpson, and Shannon indices) were calculated to assess microbial richness and evenness. Beta diversity was visualized using non-metric multidimensional scaling (NMDS) plots, generated in PRIMER version 7 (PRIMER-e, New Zealand).

Differential abundance analysis was conducted using the ANCOM-BC method via the QIIME2 q2-composition plug-in (version 2024.04). Random forest classification was used to identify key microbial taxa distinguishing between groups using MicrobiomeAnalyst 2.0 (https://www.microbiomeanalyst.ca/), with 500 decision trees and identical training/testing data sets for final model generation.

### Functional analysis

We employed the Phylogenetic Investigation of Unobserved States (PICRUSt2) algorithm to predict metabolic pathways using the MetaCyc metabolic pathway database, a comprehensive repository of experimentally validated metabolic pathways across all domains of life. To examine differences in microbial functional profiles among the WT and *Nsg2*^-/-^ groups, we applied the DESeq2 differential expression analysis algorithm. A false discovery rate (FDR)-adjusted *p*-value <0.05 and |log2FC| >1 were set to identify microbial functions that exhibited statistically significant differences between the groups. This approach enabled a systematic evaluation of the functional shifts in microbial communities associated with treatment effects.

### Statistical Analysis

Statistical analyses and visualizations were performed using GraphPad Prism 9 (GraphPad Software, USA), PRIMER version 7, and R software version 3.6.3 (R Foundation for Statistical Computing, Austria). Continuous variables are presented as mean ± SEM unless otherwise indicated. Group comparisons were performed using the Mann–Whitney U test, and correlations were assessed using Spearman’s rank correlation. A *p*-value < 0.05 was considered statistically significant. The results were then integrated with gut microbiome characteristics and metabolic pathway data to explore potential mechanistic associations.

## Results

### NSG2 expression in the colonic mucosa of a patient with ulcerative colitis

NSG2 regulates AMPA receptor trafficking in central neurons and has been identified in the developing enteric nervous system⁸, but its expression in the adult human colon and its relationship to intestinal inflammation have not been reported. We therefore examined NSG2 expression by immunohistochemistry in colonic tissue from a patient with UC and from a control subject without intestinal inflammation. NSG2 immunoreactivity was weak in the control mucosa, whereas the inflamed mucosa from the patient with UC showed markedly stronger staining, localized predominantly to the crypt epithelium (Figure 1A). This observation indicates that NSG2 is expressed in the human colon and raises the possibility that its expression increases during intestinal inflammation. Because findings from a single pair of specimens cannot establish an association between NSG2 expression and inflammation or determine the direction of causality, we next examined *Nsg2*^-/-^ mice to assess the functional effects of Nsg2 deficiency.

**Figure 1.**
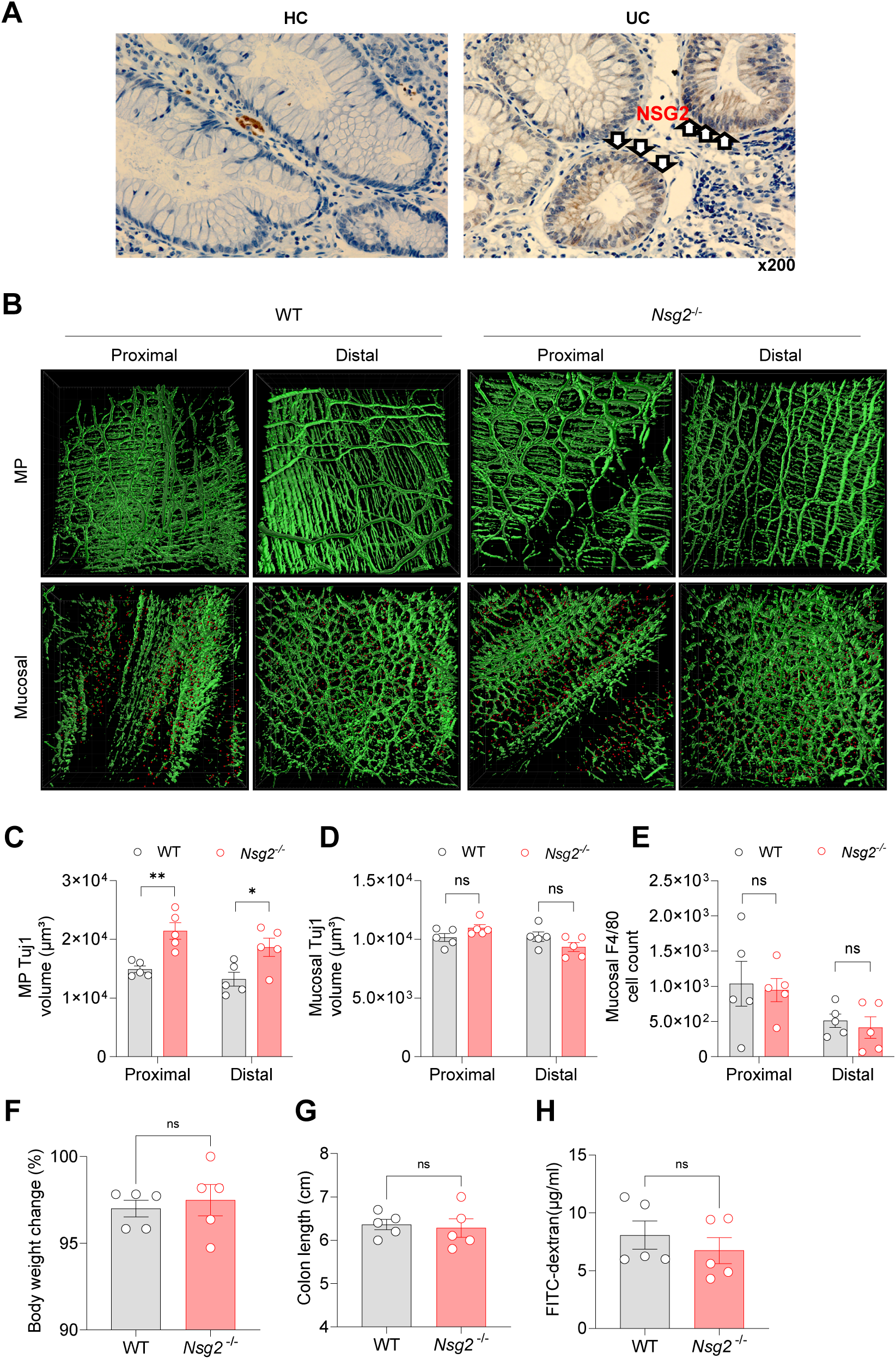
Baseline neuroimmune phenotype and intestinal permeability in WT and *Nsg2*^−/−^ mice. (A) Representative immunohistochemical staining for NSG2 in colonic mucosa from a control subject without intestinal inflammation (left) and from a patient with UC (right). Arrows indicate NSG2-immunoreactive cells. (B) Representative immunofluorescence images of colon preparations stained with Tuj1 to visualize neuronal networks in the myenteric plexus (MP, upper panels) and mucosal plexus (lower panels) from proximal and distal colon of WT and *Nsg2*^-/-^ mice. (B–C) Quantification of Tuj1^+^ nerve fiber volume in the myenteric plexus (C) and mucosal plexus (D) of proximal and distal colon. (E) Quantification of F4/80^+^ macrophages in the mucosal layer of proximal and distal colon showed no significant difference between groups. (F–H) Intestinal barrier integrity was assessed by FITC-dextran assay. No significant differences were observed in body weight change (F), colon length (G), or serum FITC-dextran concentration (H) between WT and *Nsg2*^-/-^ mice. Data are presented as mean ± SEM. Individual data points represent single mice with horizontal lines indicating group means. Statistical significance was determined by unpaired two-tailed Student’s t-test. \**p* < 0.05, \*\**p* < 0.01; ns, not significant.

### Baseline neuro and colonic immune phenotype and intestinal permeability in WT and *Nsg2*^-/-^ mice

To determine whether Nsg2 deficiency alters intestinal homeostasis before inflammatory challenge, we first characterized the steady-state intestinal phenotype of *Nsg2***^-/-^** mice. Because NSG2 regulates both synaptic development and mature synaptic transmission⁸, its constitutive deletion may affect enteric neuronal organization. Moreover, intestinal barrier integrity depends on coordinated interactions among neuronal, immune, and epithelial compartments¹³. Assessing these compartments under steady-state conditions therefore provided a baseline for interpreting subsequent differences in susceptibility to inflammation.

We first examined the ENS by quantifying Tuj1⁺ neuronal fiber density in the myenteric and mucosal plexuses of proximal and distal colon segments (Figure 1B). *Nsg2***^-/-^** mice exhibited significantly increased neuronal fiber density in the myenteric plexus of both the proximal and distal colon compared with WT controls. (Figure 1C), whereas no significant differences were observed in the mucosal plexus of either region (Figure 1D). These findings suggest that Nsg2 plays a plexus-specific role in regulating ENS organization. Given the established crosstalk between ENS and intestinal immune cells, we next assessed mucosal macrophages by F4/80 immunostaining. The abundance of F4/80⁺ macrophages were comparable between WT and *Nsg2***^-/-^**mice in both proximal and distal colons (Figure 1E), indicating that baseline immune cell populations remain unaffected by Nsg2 loss.

Given the observed changes in neuronal fiber density, we next assessed whether these structural alterations were accompanied by functional impairment of intestinal barrier integrity using FITC-dextran permeability assays. No significant differences in body weight (Figure 1F), colon length (Figure 1G), or serum FITC-dextran levels (Figure 1H) were detected between the two groups, suggesting that despite neuronal remodeling in the myenteric plexus, Nsg2 deficiency does not alter baseline intestinal barrier function under steady-state conditions.

Consistent with these findings, baseline immune profiling by flow cytometry revealed no significant differences in the frequencies of total leukocytes, CD4⁺ T cells, CD8⁺ T cells, B cells, neutrophils, monocytes, or immature and mature macrophages between WT and *Nsg2***^-/-^** mice in either the spleen (Supplementary Figure S2A) or colonic lamina propria (Supplementary Figure S2B). These results further indicate that Nsg2 deficiency does not substantially alter immune cell composition under homeostatic conditions.

Taken together, these results provide a baseline characterization of the ENS–immune– barrier axis in *Nsg2***^-/-^** mice, demonstrating that Nsg2 deficiency induces selective neuronal fiber reorganization without disrupting immune homeostasis or epithelial barrier integrity. These baseline findings prompted us to examine whether such structural neuronal alterations confer functional consequences under intestinal inflammation.

### Nsg2 deficiency confers protection against DSS-induced acute colitis

To address this question, we next employed a DSS-induced acute colitis model in WT and *Nsg2***^-/-^** mice. Mice received 3% DSS in drinking water for 6 days followed by sacrifice for analysis (Figure 2A). WT mice exhibited progressive body weight loss beginning on day 4, reaching 74.9% of initial weight by day 6 (Figure 2B). In contrast, *Nsg2*^-/-^ mice showed significantly attenuated weight loss, with significant differences observed from days 5-6 compared to WT controls. DAI, incorporating stool consistency and rectal bleeding severity, was significantly reduced in *Nsg2*^-/-^ mice compared to WT controls (*p* < 0.01, Figure 2C). Macroscopic examination revealed severe colonic inflammation with fecal blood in WT mice, while *Nsg2*^-/-^ mice displayed relatively preserved colon architecture (Figure 2D). Although colon length showed a trend toward preservation in *Nsg2*^-/-^ mice, this difference did not reach statistical significance (*p* = 0.324, Figure 2E). Histological analysis revealed extensive epithelial damage, crypt loss, and inflammatory cell infiltration in WT mice, whereas *Nsg2*^-/-^ mice maintained relatively intact mucosal architecture with minimal inflammation (Figure 2F). Correspondingly, histopathological scores were significantly lower in *Nsg2*^-/-^ mice (*p* < 0.05, Figure 2G). Together, these findings indicate that *Nsg2*^-/-^ mice are less susceptible to DSS-induced colitis, highlighting a protective role for Nsg2 deletion in acute intestinal inflammation.

**Figure 2.**
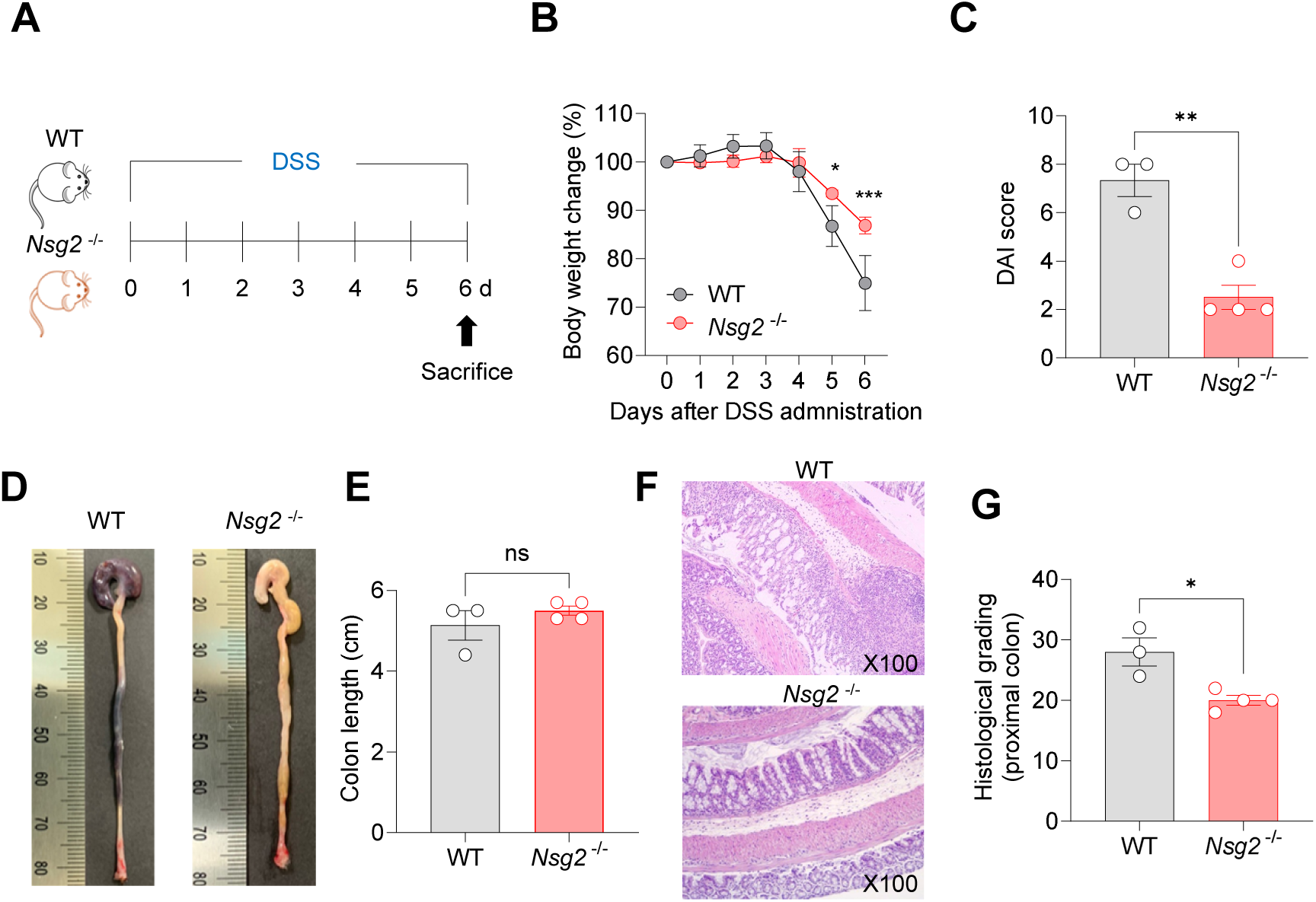
Nsg2 deficiency alleviates DSS-induced acute colitis in mice. (A) Schematic overview of the experimental design for DSS-induced colitis model. WT and *Nsg2*^-/-^ mice were administered 3% Dextran sulfate sodium (DSS) in drinking water for 6 days and were sacrificed on day 6. (B) Body weight changes monitored daily for 6 days following DSS treatment initiation (n = 4 mice/group). (C) Disease activity index (DAI) assessed on day 6. Note: one WT mouse died on day 6, reducing final no to 3. (D) Representative macroscopic images of colons harvested on day 6. (E) Colon lengths measured on day 6. (F) Representative H&E-stained sections of proximal colonic tissues collected on day 6 following DSS treatment. (G) Corresponding histological scores quantifying inflammatory damage and tissue architecture disruption. Data are presented as mean ± SEM. Individual data points represent single mice with horizontal lines indicating group means. Statistical significance was determined by unpaired two-tailed Student’s t-test. \**p* < 0.05, \*\**p* < 0.01, \*\*\**p* < 0.001; ns, not significant.

### Nsg2 deficiency suppresses colitis-associated tumorigenesis

Intestinal inflammation can promote colorectal tumorigenesis by supporting the survival and proliferation of premalignant epithelial cells. Given the reduced susceptibility of *Nsg2*^-/-^ mice to acute DSS-induced colitis, we next investigated whether Nsg2 deficiency also attenuates colitis-associated tumorigenesis using the AOM/DSS model. Mice received a single AOM injection followed by three cycles of 3% DSS treatment with recovery periods and were sacrificed on day 67 (Figure 3A). Both WT and *Nsg2^-/-^*mice maintained similar body weight trajectories throughout the protocol, indicating comparable overall health status (Figure 3B).

**Figure 3.**
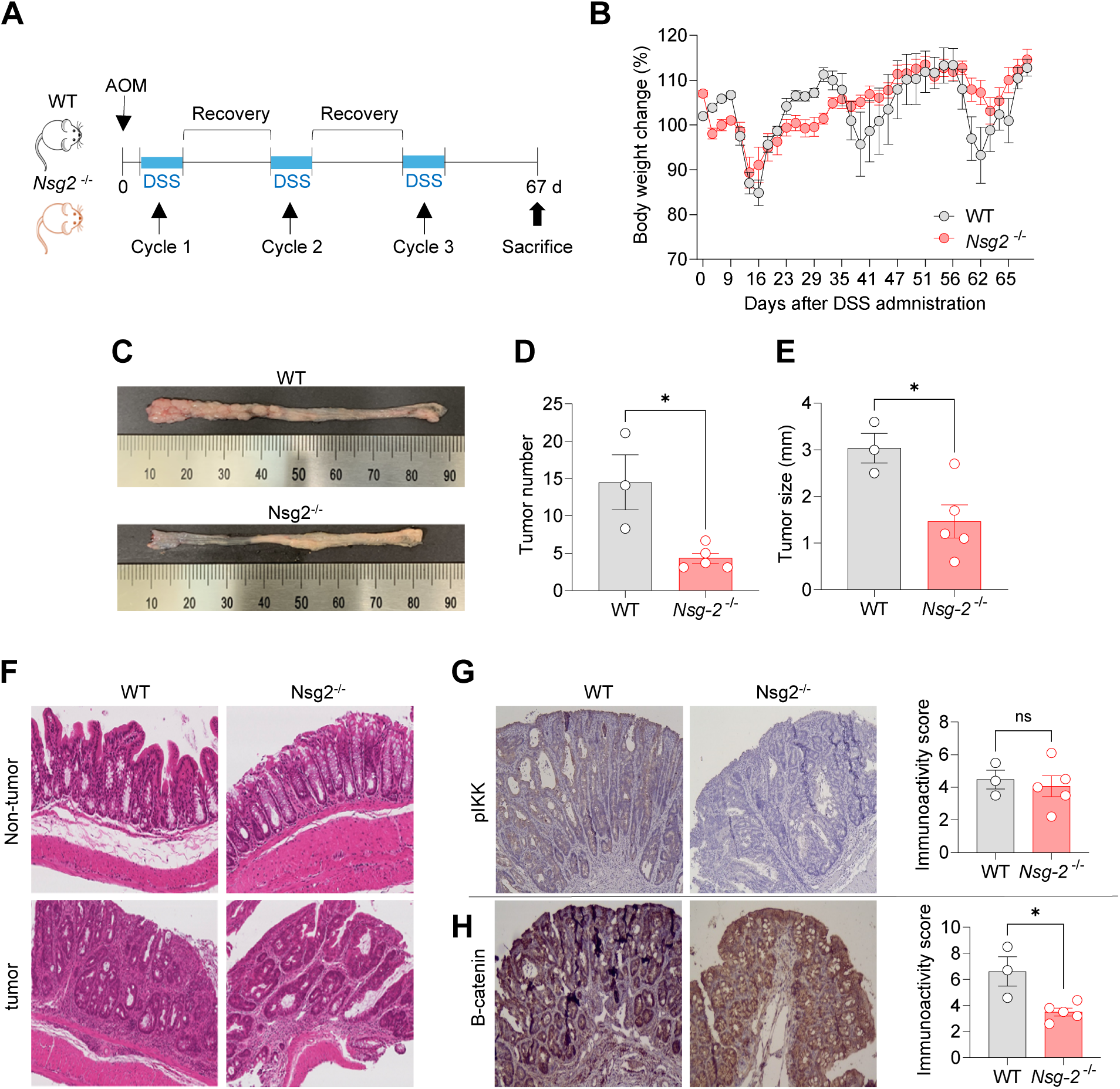
Nsg2 deficiency attenuates tumor development in the AOM/DSS-induced colitis-associated cancer model. (A) Schematic overview of the experimental design for the AOM/DSS-induced colitis-associated cancer (CAC) model. Mice received a single intraperitoneal injection of azoxymethane (AOM, 10mg/kg) followed by three cycles of 3% DSS in drinking water (5 days each), with intervening recovery periods (16 days each). Mice were sacrificed on day 67. (B) Body weight changes monitored throughout the AOM/DSS protocol (n = 5 mice/group initially; two WT mice died on day 63, final n =3 for WT). (C) Representative macroscopic images of the colons analyzed on day 67. (D) Quantification of tumor size and (E) number in WT and *Nsg2*^-/-^ mice. (F) Representative H&E-stained sections of distal colon showing tumor and adjacent non-tumor areas collected on day 67. (G) Representative immunohistochemical staining for phosphorylated IkB kinase (pIKK) in tumor tissues with immunoreactivity scores (right panel). (H) Representative immunohistochemical staining for β-catenin (H) in tumor tissues with quantification of immunoactivity scores (right panel). Data are presented as mean ± SEM. Individual data points represent single mice with horizontal lines indicating group means. Statistical significance was determined by unpaired two-tailed Student’s t-test. \**p* < 0.05; ns, not significant.

However, macroscopic examination revealed markedly reduced tumor burden in *Nsg2^-/-^* mice compared to WT controls (Figure 3C). Quantitative analysis demonstrated that *Nsg2^-/-^*mice developed significantly fewer tumors (*p* < 0.05, Figure 3D) and smaller tumor sizes (*p* < 0.05, Figure 3E) compared to WT mice. Histological analysis revealed disrupted epithelial architecture and dense inflammatory infiltration in WT tumors, whereas tumors from *Nsg2^-/-^*mice showed relatively preserved mucosal architecture (Figure 3F). To further investigate potential mechanisms, we examined key signaling pathways by immunohistochemistry. pIKK expression showed no significant difference between genotypes (Figure 3G). In contrast, β-catenin expression was significantly reduced in *Nsg2^-/-^*tumors compared to WT tumors (*p* < 0.05, Figure 3H), suggesting altered Wnt signaling. These findings demonstrate that Nsg2 deficiency attenuates colitis-associated tumor development, potentially through modulation of Wnt/β-catenin signaling, identifying Nsg2 as a pro-tumorigenic factor in inflammation-driven colorectal cancer.

### Nsg2 deficiency enhances macrophage activation-associated features without altering polarization

To investigate whether the protective effects of Nsg2 deficiency involve alterations in immune cell recruitment or activation, we performed a comprehensive flow cytometric analysis of colonic lamina propria following AOM/DSS treatment. Quantitative analysis of major immune cell populations using cell gating strategy (Supplementary Figure S1) revealed no significant differences in the frequency of total leukocytes, CD4^+^ T cells, CD8^+^ T cells, B cells, or neutrophils between WT and *Nsg2*^-/-^ mice (Supplementary Figure S3A). Similarly, myeloid cell subsets, including monocytes, immature macrophages, and mature macrophages, showed comparable abundance between genotypes, indicating that Nsg2 deficiency does not grossly alter innate immune cell recruitment to the inflamed colon. High-dimensional analysis using t-distributed Stochastic Neighbor Embedding (t-SNE) confirmed these findings, revealing similar overall distribution patterns and clustering of CD45^+^ immune cells (Supplementary Figure S3B), myeloid cells, T cells, and B cells (Supplementary Figure S3C) between WT and *Nsg2*^-/-^ mice. These data demonstrate that the protective phenotype observed in *Nsg2*^-/-^ mice is independent of major changes in immune cell composition within the tumor microenvironment.

Since macrophages represent a dominant innate immune population in the inflamed colon and are key regulators of tumor-promoting inflammation,^12,13^ we next focused on macrophage polarization states to determine whether Nsg2 deficiency selectively reprograms macrophage function. Macrophage subsets were identified by gating CD45⁺CD11b⁺Ly6G⁻CD64⁺ cells (Supplementary Figure S4A). Quantitative analysis revealed comparable frequencies of M1 macrophages (MHCII^high^CD206^low^) and M2 macrophages (MHCII^high^CD206^high^) between WT and *Nsg2*^-/-^ mice, with no significant difference in the M1/M2 ratio (Supplementary Figure S4B), indicating that Nsg2 deficiency does not alter macrophage polarization under inflammatory conditions.

Despite the absence of polarization shifts, CD86 and IL-6 expression levels were significantly elevated in macrophages from *Nsg2*^-/-^ mice compared with WT controls (p < 0.05; Supplementary Figure S4C–E). In contrast, M2-associated functional markers, including CD206 and IL-10, showed no significant changes in *Nsg2*^-/-^ mice relative to WT controls (Supplementary Figure S4F). This enhanced activation signature, paradoxically occurring alongside reduced tumor burden and attenuated inflammation, may reflect a context-dependent anti-tumor macrophage response, the precise mechanisms of which require further investigation.

Taken together, these results demonstrate that Nsg2 deficiency does not substantially alter immune cell composition or macrophage polarization but is associated with increased expression of macrophage activation-related markers. Thus, the protective phenotype observed in *Nsg2*^-/-^ mice is unlikely to be driven by broad changes in immune cell recruitment or macrophage polarization, supporting the involvement of alternative mechanisms.

### Co-housing eliminates the protective effects of Nsg2 deficiency

To determine whether the protective effects of Nsg2 deficiency are microbiome-dependent, we co-housed WT and *Nsg2*^-/-^ mice for 6 weeks prior to experimental treatments to equilibrate their gut microbiota. In the DSS-induced colitis model (Figures 4A), co-housed mice showed similar disease progression regardless of genotype. Both WT and *Nsg2*^-/-^ mice exhibited comparable body weight loss (Figure 4B), DAI scores (Figure 4C), and macroscopic colon appearance (Figure 4D). Colon length measurements revealed no significant differences between groups (Figure 4E). Histological analysis showed similar degrees of epithelial damage and inflammatory infiltration in both genotypes (Figure 4F), with comparable histopathological scores (Figure 4G). In the AOM/DSS tumor model (Figures 5A), co-housed mice maintained similar body weights throughout the protocol (Figures 5B). Macroscopic examination revealed comparable colon morphology between groups (Figure 5C), with no differences in colon length (Figure 5D). Importantly, tumor burden analysis showed no significant differences in either tumor number (Figure 5E) or tumor size (Figure 5F) between co-housed WT and *Nsg2*^-/-^ mice. These results demonstrate that co-housing eliminates both the protective effect against colitis and the tumor-suppressive effect observed in separately housed *Nsg2*^-/-^ mice. These findings suggest that alterations in the gut microbiome contribute substantially to the protective effects associated with Nsg2 deficiency.

**Figure 4.**
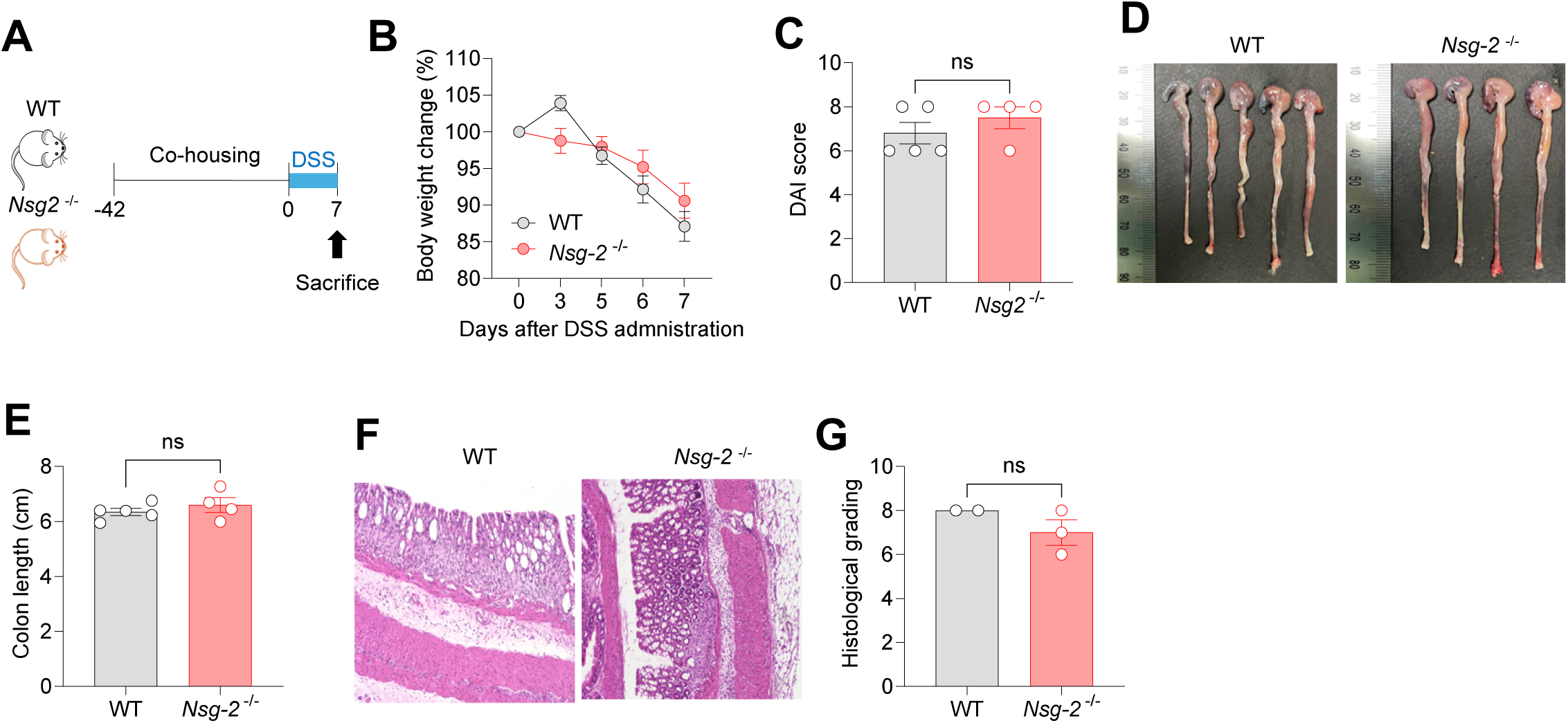
Cohousing modulates susceptibility to colitis in *Nsg2*-deficient mice. (A) Schematic overview of the experimental design. WT and *Nsg2*^-/-^ mice were co-housed for 6 weeks prior to DSS treatment to allow microbiome equilibration. Mice were subsequently administered 3% DSS in drinking water for 7 days and sacrificed on day 7. (B) Body weight changes monitored daily for 7 days following DSS treatment initiation (WT, n = 5; *Nsg2*^-/-^, n = 4). (C) Disease activity index (DAI) assessed on day 7. (D) Macroscopic images of colons harvested on day 7. (E) Colon lengths measured on day 7. (F) Representative H&E-stained sections of distal colonic tissues collected on day 7 following DSS treatment. (G) Corresponding histological scores quantifying inflammatory damage and tissue architecture disruption. Data are presented as mean ± SEM; Individual data points represent single mice with horizontal lines indicating group means. Statistical significance was determined by unpaired two-tailed Student’s t-test. ns, not significant.

**Figure 5.**
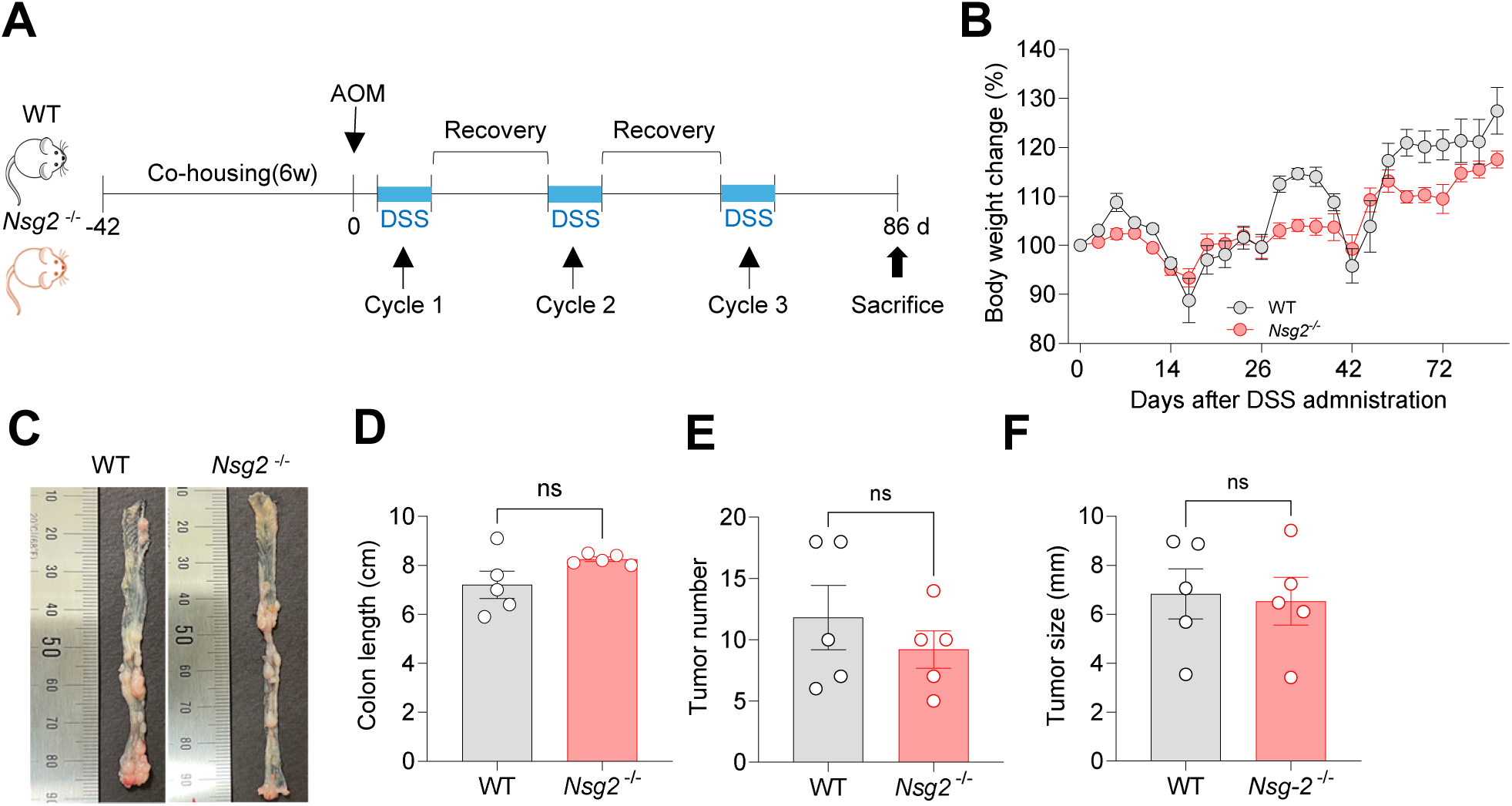
Cohousing modulates susceptibility to CAC in *Nsg2*-deficient mice. (A) Schematic overview of the experimental design. WT and *Nsg2*^-/-^ mice were co-housed for 6 weeks prior to DSS treatment to allow microbiome equilibration. Mice then received a single intraperitoneal injection of azoxymethane (AOM, 10 mg/kg) followed by three cycles of 3% DSS in drinking water (5 days each), with intervening recovery periods (16 days each). Mice were sacrificed on day 86. (B) Body weight changes monitored throughout the AOM/DSS protocol (n = 5 mice/group). (C) Representative macroscopic images of colons analyzed on day 86. (D) Colon lengths measured on day 86. (E) Quantification of tumor number and (F) tumor size in co-housed WT and *Nsg2*^-/-^ mice. Data are presented as mean ± SEM. Individual data points represent single mice with horizontal lines indicating group means. Statistical significance was determined by unpaired two-tailed Student’s t-test. ns, not significant.

### Nsg2 deficiency alters gut microbiome composition and diversity

To investigate whether the protective effects of Nsg2 deficiency are associated with changes in gut microbiota, we performed 16S rRNA gene sequencing (V3-V4 region) on fecal samples from WT, *Nsg2*^-/-^, co-housed WT, and co-housed *Nsg2*^-/-^ mice. Taxonomic composition analysis revealed distinct microbial profiles between genotypes, with *Nsg2*^-/-^ mice showing altered abundance of several bacterial genera compared to WT controls (Figure 6A). Alpha-diversity analysis demonstrated significantly greater microbial richness (observed features) and diversity (Shannon and Simpson indices) in Nsg2^-/-^ mice than in WT mice (Figure 6B). Notably, co-housed mice displayed intermediate alpha-diversity levels between their respective genotype groups, although the differences were not statistically significant (*p* > 0.05), suggesting partial microbiota convergence through environmental sharing (Figure 6C). Beta-diversity analysis revealed distinct microbial community clustering patterns, with *Nsg2*^-/-^ mice separating significantly from WT controls (Figure 6, *p* < 0.05). Co-housed mice clustered intermediately between their genotype groups, confirming the impact of shared housing on microbial composition. To identify specific taxa driving these differences, we performed differential abundance analysis using ANCOM-BC on non-co-housed samples. This analysis revealed significant log fold changes in multiple bacterial features (Figure 6D). The top genera enriched in *Nsg2^-/-^* mice (red bars) included *Butyricicoccus*, *Ruminococcaceae*, *ASF356*, *Helicobacter*, and *Lachnospiraceae*_NK4A136_group, while those enriched in WT mice (blue bars) included *Faecalibaculum*, *Bifidobacterium*, *Enterorhabdus, Rikenellaceae_RC9_gut_group*, and *Enterococcus*. Random forest machine learning analysis further confirmed these discriminatory taxa, with *RF39*, *Faecalibaculum, Turicibacter, Eubacterium_nodatum_group and ASF356* among the top features distinguishing between genotypes (Figure 6E). Detailed examination of individual genera confirmed that *Faecalibaculum*, *Bifidobacterium*, *Enterorhabdus, RF39,* and Turicibacter were significantly more abundant in WT mice (Figure 6F). Conversely, genera including *Butyricicoccus*, *Ruminococcaceae*, *ASF356*, *Helicobacter*, and *Lachnospiraceae_NK4A136_group* were significantly more abundant in *Nsg2^-/-^* mice (Figure 6G). These results demonstrate that Nsg2 deficiency leads to substantial remodeling of the gut microbiome, characterized by increased microbial diversity and specific taxonomic shifts that may contribute to the observed protective phenotypes against colitis and tumorigenesis.

**Figure 6.**
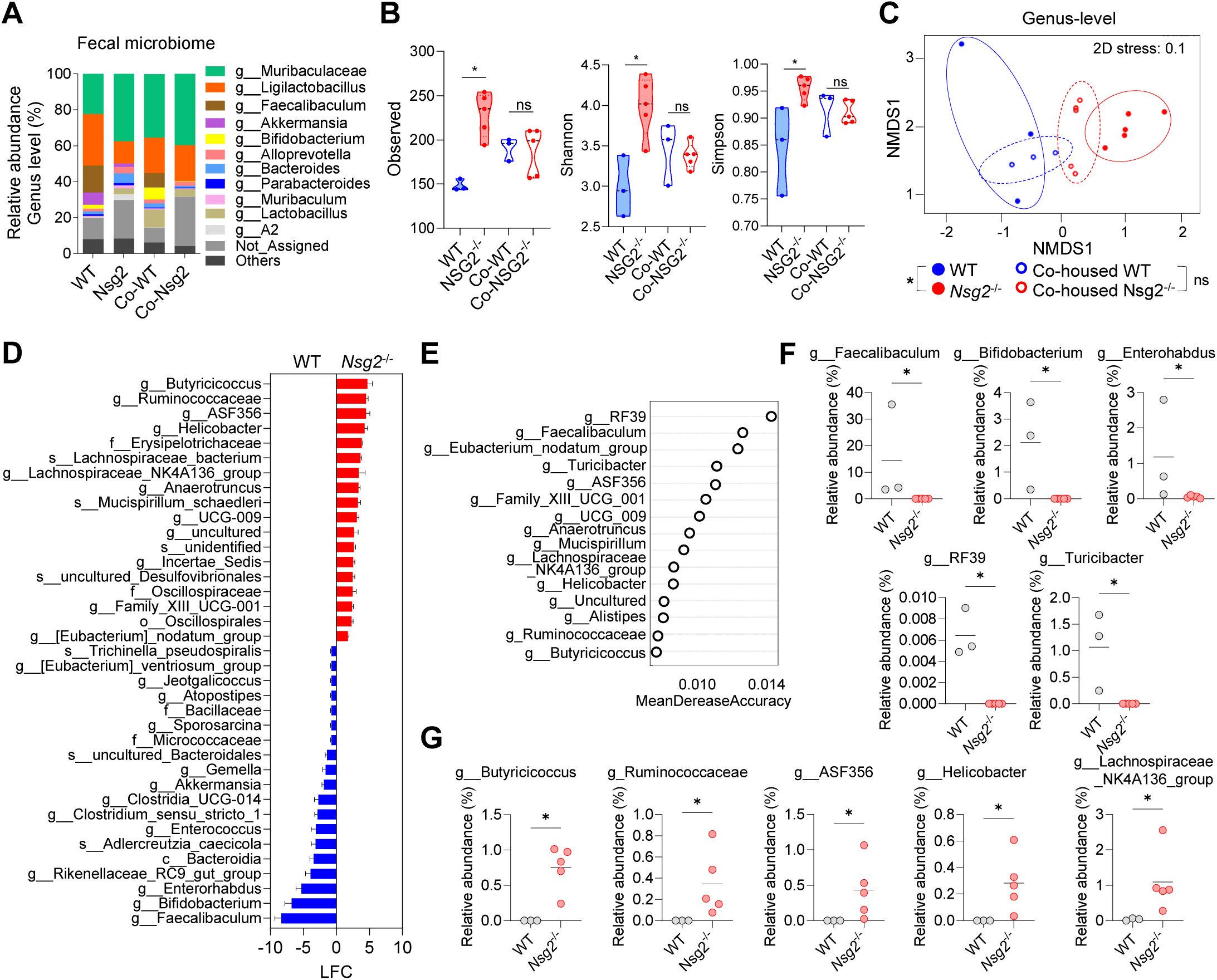
Nsg2 deficiency alters the gut microbiome. (A-C) Fecal microbiome analysis of WT (n=3), *Nsg2*^-/-^ (n=5), co-housed WT (n=3), co-housed *Nsg2*^-/-^ (n=5) mice by 16S rRNA amplicon sequencing. (A) Relative abundance of fecal bacteria at the genus level. (B) Alpha-diversity indices including Observed features, Simpson’s index, and Shannon’s diversity index. (C) Non-metric multidimensional scaling (NMDS) plots at amplicon sequence variant (ASVs) level for beta-diversity. (D-G) Differential abundance analysis between non-co-housed WT and *Nsg2*^-/-^ mice. (D) ANCOM-BC (Analysis of Compositions of Microbiomes with Bias Correction) analysis at the feature level showing log fold change (LFC) of bacterial features between groups. (E) Random Forest analysis at the genus level showing mean decrease accuracy for discriminative taxa. (F) Relative abundance of representative genera significantly enriched in WT mice, selected based on results from panels D and E. (G) Relative abundance of representative genera significantly enriched in *Nsg2*^-/-^ mice. Data in panel (B) are presented as violin plots showing the full distribution (minimum to maximum) with median indicated by the central line. *p*-values in panel (C) were determined by PERMANOVA (Permutational Multivariate Analysis of Variance) Benjamini–Hochberg FDR correction. Individual data points in panels (F) and (G) represent individual mice, with horizontal lines indicating group means. \**p* < 0.05; ns, not significant.

### Nsg2 deficiency alters the functional potential of the gut microbiome

To investigate the functional consequences of the taxonomic changes observed in *Nsg2^-/-^* mice, we performed predictive functional profiling using PICRUSt2 analysis based on 16S rRNA sequencing data. COG (Clusters of Orthologous Groups) functional classification revealed distinct metabolic profiles between WT and *Nsg2*^-/-^ mice across 22 functional categories (Figure 7A). The analysis showed that metabolic functions comprised the largest proportion of predicted microbial capabilities, including amino acid transport and metabolism, carbohydrate transport and metabolism, and coenzyme transport and metabolism. Cellular processes and signaling represented the second most abundant category (23%), followed by information storage and processing (17%). Notably, *Nsg2^-/-^* mice exhibited enhanced representation in several metabolic categories, particularly those related to energy production and conversion. LEfSe (Linear Discriminant Analysis Effect Size) analysis identified specific KEGG orthologs that were differentially abundant between genotypes (Figure 7B). Key functional genes enriched in *Nsg2^-/-^* mice (red bars) included neurotransmitter:Na+ symporter, CAAX protease family proteins, U32 family peptidases, cobyrinic acid a,c-diamide synthetase, and TatD DNase family proteins, reflecting enhanced capabilities in membrane transport, protein processing, and DNA repair mechanisms. Conversely, WT mice (gray bars) showed enrichment of ABC-2 type transport system ATP-binding proteins, 3-oxoacyl reductases, aldehyde:ferredoxin oxidoreductases, reverse gyrase, glutamate synthase, and multicomponent Na+:H+ antiporter subunits. Random forest analysis further validated these functional differences, identifying discriminatory KO features with high predictive accuracy (Figure 7C). Top-ranking features enriched in *Nsg2^-/-^*mice (red circles) included methenyltetrahydromethanopterin cyclohydrolase (K01499), ribosomal protein S12 methylthiotransferase accessory factor (K09136), queuosine precursor transporter (K09125), and tryptamine:L-glutamate ligase (K06914). These enzymes are primarily involved in one-carbon metabolism, oxidative stress resistance, and amino acid metabolism, suggesting enhanced metabolic flexibility in *Nsg2^-/-^* microbiota. KEGG pathway enrichment analysis revealed significant upregulation of specific metabolic pathways in *Nsg2^-/-^*mice (Figure 7D). Most notably, methane metabolism showed the highest enrichment (*p* = 2.57 × 10^-11^), followed by porphyrin and chlorophyll metabolism (*p* = 2.03 × 10^-5^), and one-carbon pool by folate metabolism (*p* < 0.05). Additional pathways showing significant enrichment included pyruvate metabolism, biotin metabolism, cysteine and methionine metabolism, and glutathione metabolism (all *p* < 0.05), indicating broad metabolic reprogramming in the *Nsg2^-/-^* microbiome.

**Figure 7.**
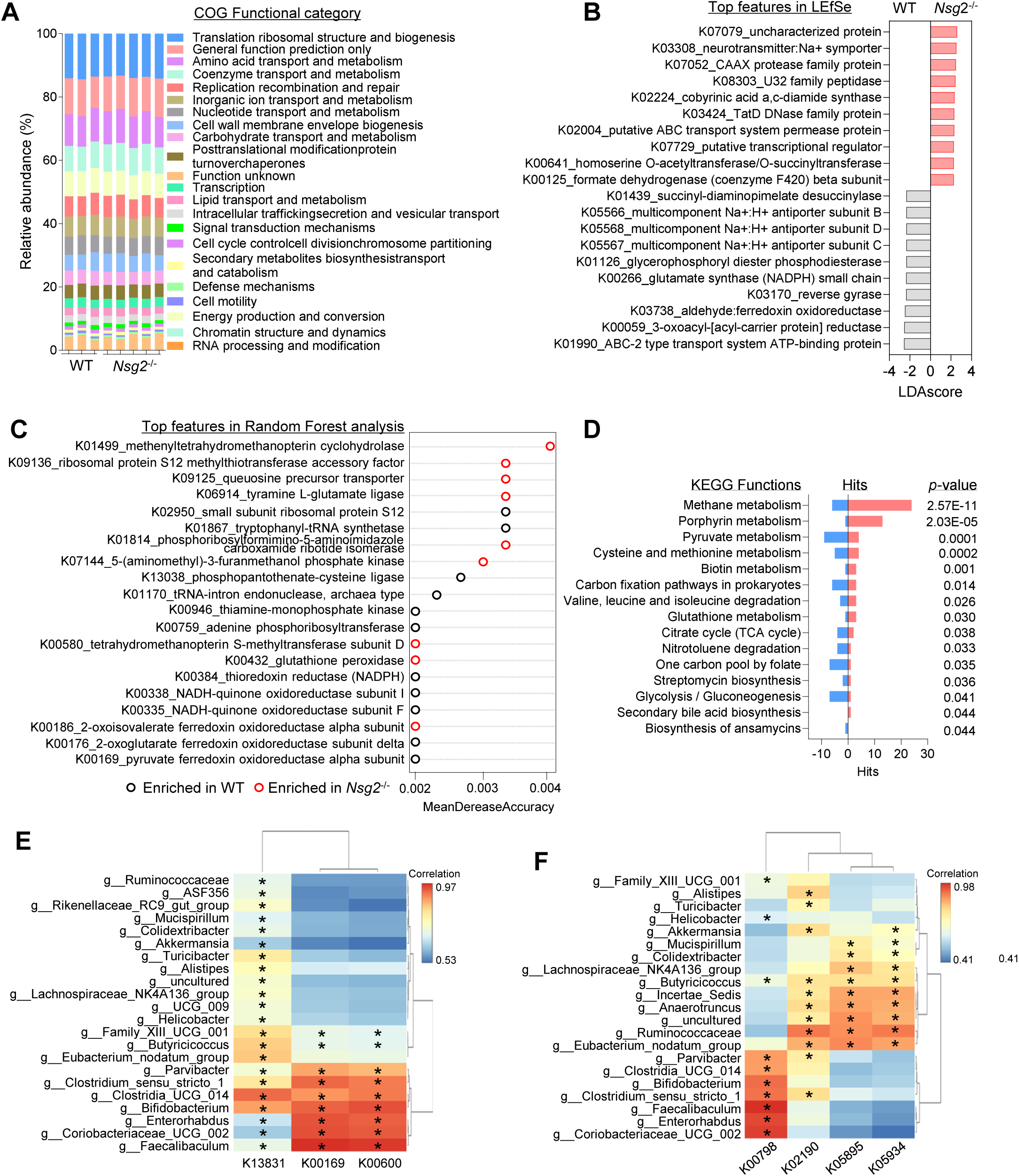
Functional profiling of gut microbiota in WT and *Nsg2*-deficient mice based on PICRUSt2 prediction. (A) Relative abundance of predicted COG functional categories in WT and *Nsg2*^-/-^ mice, showing differences in metabolic, signaling, and cellular stress response functions. (B) LEfSe (Linear discriminant analysis Effect Size) analysis highlighting significantly differentially abundant KEGG orthologs (KOs) between WT and *Nsg2*^-/-^ mice. Gray and red bars represent features significantly enriched in WT and *Nsg2*^-/-^ mice, respectively. (C) Random forest analysis identifying the top discriminative KO features between WT and *Nsg2*^-/-^ mice. Blue-labeled KOs are enriched in WT mice, while red-labeled KOs are enriched in *Nsg2*^-/-^ mice. (D) KEGG pathway enrichment analysis showing altered functional pathways between WT and *Nsg2*^-/-^ mice, with notable increases in methane metabolism and porphyrin biosynthesis pathways. (E) Heatmap showing distance correlation analysis between methane metabolism-related genes and enriched bacterial taxa at the genus level. Key enzymes include: K13831 (hps-phi, 3-hexulose-6-phosphate synthase/6-phospho-3-hexuloisomerase); K00169 (porA, pyruvate ferredoxin oxidoreductase alpha subunit); K00600 (glyA, glycine hydroxymethyltransferase) (F) Heatmap showing distance correlation analysis between porphyrin metabolism-related genes and bacterial taxa at the genus level. Key enzymes include: K00798 (MMAB, cob(I)alamin adenosyltransferase); K20190 (cbiK, sirohydrochlorin cobaltochelatase); K05895 (precorrin-6A/cobalt-precorrin-6A reductase); K05934 (precorrin-3B C17-methyltransferase/cobalt-factor III methyltransferase). Asterisks in panels (E) and (F) indicate statistically significant correlations (adjusted *p* < 0.05).

To explore the relationship between specific bacterial taxa and these metabolic functions, we performed distance correlation analysis. For methane metabolism, key enzymes including 3-hexulose-6-phosphate synthase (K13831), pyruvate ferredoxin oxidoreductase (K00169), and glycine hydroxymethyltransferase (K00600), showed significant correlations with several bacterial genera enriched in *Nsg2^-/-^* mice (Figure 7E). Similarly, porphyrin metabolism-related enzymes, including cob(I)alamin adenosyltransferase (K00798), sirohydrochlorin cobaltochelatase (K20190), precorrin-6A reductase (K05895), and precorrin-3B methyltransferase (K05934), exhibited significant correlations with specific taxa (Figure 7F). These results demonstrate that Nsg2 deficiency not only alters gut microbiome composition but also significantly enhances the functional potential of the microbial community, particularly in pathways related to methane metabolism, porphyrin biosynthesis, and one-carbon metabolism. This functional reprogramming may contribute to the protective effects observed against intestinal inflammation and tumorigenesis in *Nsg2^-/-^* mice.

## Discussion

This study provides novel insights into the role of Nsg2 in intestinal inflammation and tumorigenesis, identifying it as a key modulator within the ENS-microbiome axis. Using DSS-induced acute colitis and CAC models, we demonstrated that Nsg2 deficiency significantly alleviates intestinal inflammation, preserves epithelial integrity, and attenuates tumor development. These protective phenotypes were accompanied by marked alterations in gut microbial community composition and predicted functional capacity. Co-housing experiments further revealed that the beneficial effects of Nsg2 deficiency are largely microbiome-dependent, suggesting a mechanistic link between Nsg2 signaling, gut microbial ecology, and host immune regulation. Predictive metagenomic analysis indicated that Nsg2 deletion is associated with enrichment of beneficial bacterial genera and upregulation of microbial metabolic pathways, particularly methane and one-carbon metabolism, which may contribute to intestinal homeostasis and protection against inflammation-driven carcinogenesis.

The marked attenuation of DSS-induced colitis in *Nsg2*^-/-^ mice suggests a previously unrecognized role for Nsg2 in regulating intestinal inflammation. The ENS has emerged as a critical regulator of mucosal immunity, with ENS-derived neurotransmitters capable of shaping macrophage phenotype and T cell function within the lamina propria.^14,15^ Given Nsg2’s role in excitatory synaptic transmission, its deletion may attenuate aberrant neuronal signaling under inflammatory conditions, thereby dampening neurogenic inflammation. However, this hypothesis remains to be directly tested. Notably, immune profiling of the colonic lamina propria revealed that while M1/M2 macrophage polarization ratios were comparable between genotypes, the expression of activation-associated markers CD86 and IL-6 was paradoxically elevated in *Nsg2*^-/-^ mice. This dissociation between activation state and polarization balance, occurring in the context of reduced histological inflammation and epithelial damage, suggests that Nsg2 may regulate intestinal inflammation through noncanonical pathways beyond classical macrophage polarization. Whether this reflects Nsg2-mediated neuroimmune crosstalk or independent immune reprogramming warrants further investigation.

Beyond microbial compositional changes, we sought to identify the downstream oncogenic signaling pathways underlying the reduced tumor burden in *Nsg2*^-/-^ mice. In the AOM/DSS-induced CAC model, *Nsg2*^-/-^ mice exhibited a significantly reduced tumor burden, characterized by both decreased tumor number and size relative to WT controls. Mechanistic investigations revealed a marked reduction in β-catenin expression and nuclear translocation in tumors from *Nsg2*^-/-^ mice (Figure 3H), indicative of suppressed Wnt/β-catenin signaling, a key oncogenic pathway in colorectal tumorigenesis. In contrast, expression of phosphorylated IκB kinase (pIKK), a canonical marker of NF-κB pathway activation, did not differ significantly between genotypes (Figure 3G). These results suggest that Nsg2 may promote tumor progression predominantly through modulation of Wnt/β-catenin-driven transcriptional programs, rather than through NF-κB signaling. This mechanistic insight aligns with emerging evidence that neuronal activity can contribute to tumor development via neurotransmitter-mediated signaling within the tumor microenvironment, potentially including modulation of Wnt-dependent transcriptional programs.^16–18^

In support of a microbiome-mediated mechanism, our co-housing experiments demonstrated that the protective effects of Nsg2 deficiency were largely abolished when *Nsg2*^-/-^ mice shared the same microbial environment with WT mice.^19^ Co-housed *Nsg2*^-/-^ mice exhibited similar disease severity, histological inflammation, and tumor burden as co-housed WT mice in both colitis and CAC models, underscoring the central role of gut microbial composition in mediating the Nsg2-dependent protective phenotype.^20^ 16S rRNA sequencing revealed that Nsg2 deficiency was associated with increased microbial diversity and a distinct shift in bacterial community structure, including enrichment of potentially beneficial taxa such as Butyricicoccus and members of the Lachnospiraceae family and depletion of potentially pro-inflammatory genera including Faecalibaculum, Enterorhabdus, Bifidobacterium, RF39, Turicibater. These compositional changes were accompanied by functional reprogramming of the microbiota, as predictive metagenomic analysis identified significant enrichment of metabolic pathways related to methane metabolism, porphyrin biosynthesis, and one-carbon metabolism in *Nsg2*^-/-^ mice. Such pathways are increasingly recognized for their roles in regulating intestinal barrier integrity, immune tolerance, and oxidative stress, all of which are essential for maintaining gut homeostasis.^21,22^ Collectively, these findings suggest that Nsg2 modulates gut inflammation and tumorigenesis at least in part through reshaping the metabolic and taxonomic landscape of the intestinal microbiome.^23,24^

Interestingly, several bacterial taxa altered in *Nsg2*^-/-^ mice, including members of the Ruminococcaceae and Lachnospiraceae families, overlap with microbial signatures reported in neuromodulatory gut disorders such as IBS.^25,26^ Notably, these taxa are typically enriched in healthy individuals but are significantly depleted in patients with IBS, consistent with their association with gut homeostasis and anti-inflammatory functions. In contrast, we observed a relative enrichment of these bacterial groups in Nsg2-deficient mice, suggesting a shift toward a microbiota configuration associated with a healthier gut state. While direct comparisons between murine and human microbiota require caution, these opposing trends support the notion that Nsg2 deficiency may promote beneficial microbial remodeling, potentially extending beyond IBD-associated inflammation to influence broader patterns of gut dysbiosis. Given the established contribution of the ENS to both IBD and IBS pathophysiology^27^, our findings support the hypothesis that Nsg2 may modulate microbiota–ENS–immune interactions relevant to intestinal homeostasis and disease susceptibility, though validation in human cohorts will be necessary to substantiate this broader implication.

In *Nsg2*^-/-^ mice, functional predictions revealed significant upregulation of microbial pathways involved in methane and porphyrin metabolism. Methane-related microbial functions have been associated with altered intestinal physiology and microbial metabolic interactions.^28–30^ In the present study, the enrichment of predicted methane-related pathways in Nsg2⁻/⁻ mice suggests a shift in microbial functional potential rather than direct evidence of increased methane production. Notably, enrichment of K13831 (3-hexulose-6-phosphate synthase), a key enzyme in C1 carbon assimilation, was positively correlated with *Nsg2*^-/-^-enriched taxa including Butyricicoccus and Eubacterium nodatum group, suggesting an enhanced capacity for formaldehyde detoxification and cytoprotective metabolite production.^31–33^

Microbial porphyrin metabolism was also differentially enriched, with three biosynthetic enzymes (K20190, K05895, and K05934) positively correlated with taxa enriched in *Nsg2*^-/-^ mice. This shift in porphyrin biosynthesis, combined with enhanced C1 metabolism, suggests functional reprogramming of the microbiome toward improved redox balance, epithelial integrity, and immune regulation.^34,35^ In contrast, K00798 (cob(I) alamin adenosyltransferase), associated with conventional cobalamin-dependent metabolism, was enriched in WT mice, further highlighting the distinct metabolic landscape of *Nsg2*^-/-^ microbiota. Collectively, the concurrent enrichment of methane and porphyrin metabolic pathways in *Nsg2*^-/-^ mice suggests a coordinated functional reprogramming of the gut microbial community toward anti-inflammatory and cytoprotective states, which may underlie the observed attenuation of colitis and tumorigenesis.

Several limitations of this study warrant acknowledgment. First, the absence of littermate controls raises the possibility of confounding genetic drift effects. Second, while co-housing experiments provided strong evidence for microbiome-dependent mechanisms, definitive validation would benefit from complementary approaches such as fecal microbiota transplantation or antibiotic depletion studies. Third, functional pathway predictions were derived from PICRUSt2, an inference-based tool, and thus require validation through direct metagenomic or metabolomic approaches. Fourth, the precise molecular interactions between Nsg2, neurotransmitter receptors, and immune cell trafficking pathways require further mechanistic elucidation. Finally, the clinical relevance of these findings awaits validation in human IBD and colorectal cancer contexts.

In conclusion, this study identifies Nsg2 as a novel modulator of intestinal inflammation and tumorigenesis, functioning at the intersection of neuronal signaling, immune regulation, and microbial ecology. Nsg2 deficiency conferred protection against DSS-induced colitis and reduced tumor development in the AOM/DSS model, with protective effects that were largely microbiome-dependent and associated with suppression of Wnt/β-catenin oncogenic signaling. These findings suggest that Nsg2 may represent a potential therapeutic target for IBD and inflammation-driven colorectal cancer, pending further validation. Future investigations should focus on elucidating the precise molecular interactions between Nsg2 and gut microbial communities, validating functional metabolic pathways through direct metagenomic and metabolomic approaches, and ultimately translating these findings to human disease contexts through clinical studies.

## Author contributions

G.K., K.Y.C., H.S.Y., and K.J.C. performed the experiments. G.K. and K.Y.C. analyzed the data. S.K. supervised the project. G.K., K.Y.C., and S.K. wrote the manuscript with support from H.P, J.P.L., B.G.K., J.S.K. All authors had access to the study data and had reviewed and approved the final manuscript.

---

**Supplementary Figure S1.**
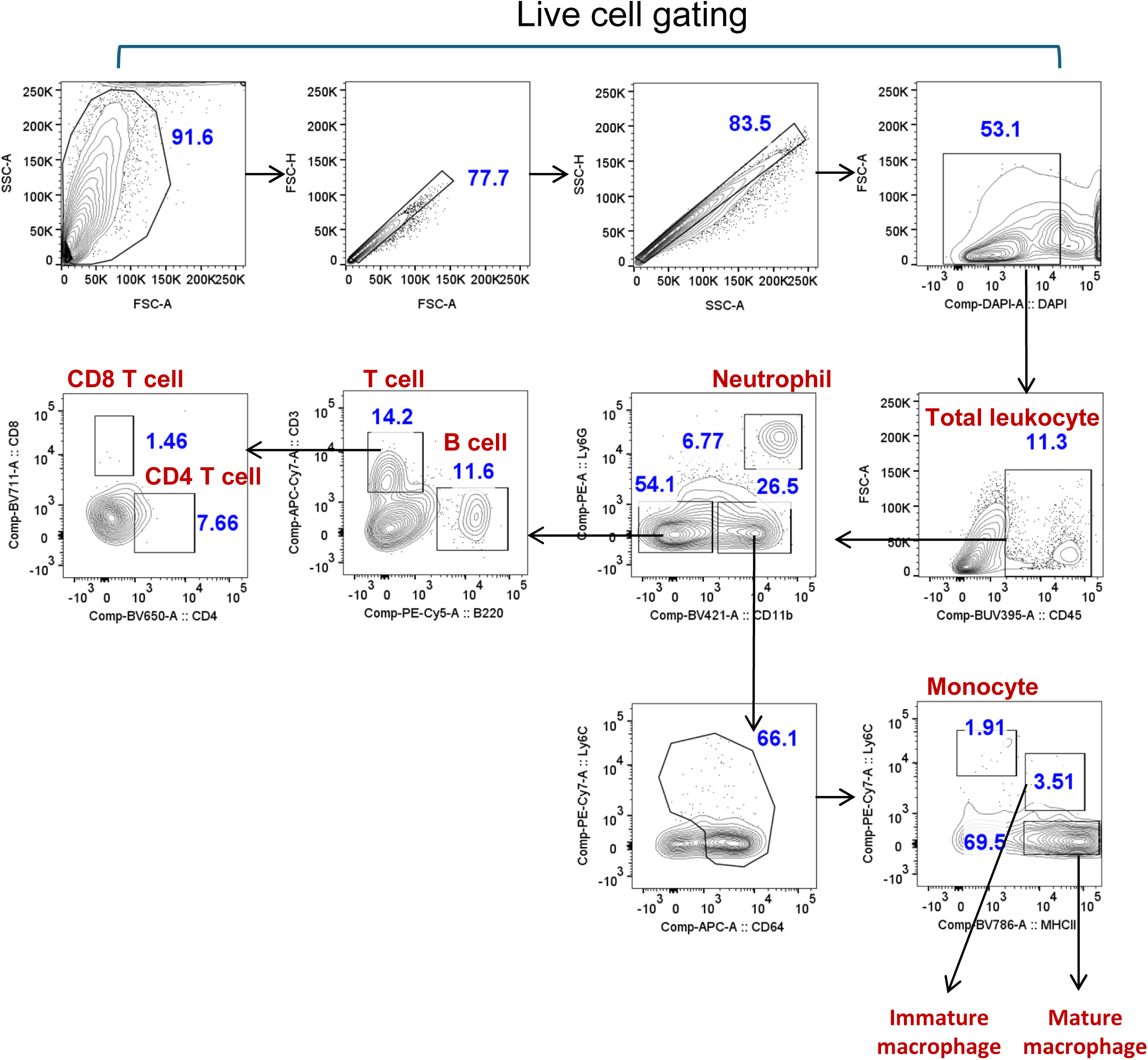
Flow cytometry gating strategy for immune cell profiling. Representative flow cytometry plots showing the sequential gating strategy used for identification and quantification of major immune cell populations in colonic lamina propria. Single cells were first identified using forward scatter area (FSC-A) vs. forward scatter height (FSC-H) to exclude doublets, followed by side scatter area (SSC-A) vs. FSC-A gating to select viable cells based on morphological parameters. Live cells were distinguished from dead cells using a viability dye. From the live cell population, major immune cell subsets were identified as follows: total leukocytes (CD45^+^), CD8^+^ T cells (CD3^+^CD8^+^), CD4^+^ T cells (CD3^+^CD4^+^), B cells (CD3^-^B220^+^), neutrophils (CD11b^+^Ly6G^+^) and monocyte/macrophage populations (CD11b^+^CD64^+^). Monocytes and macrophages were further distinguished based on Ly6C expression, with Ly6C^high^ cells representing inflammatory monocytes/immature macrophages and Ly6C^low^ cells representing mature tissue-resident macrophages. Numbers in each gate represent the percentage of parent population. This gating strategy was consistently applied across all experimental groups to ensure accurate quantification of immune cell populations in both WT and *Nsg2*^-/-^ mice following AOM/DSS treatment. Analysis was performed using FlowJo software (Tree Star Inc., Ashland, OR, USA).

**Supplementary Figure S2.**
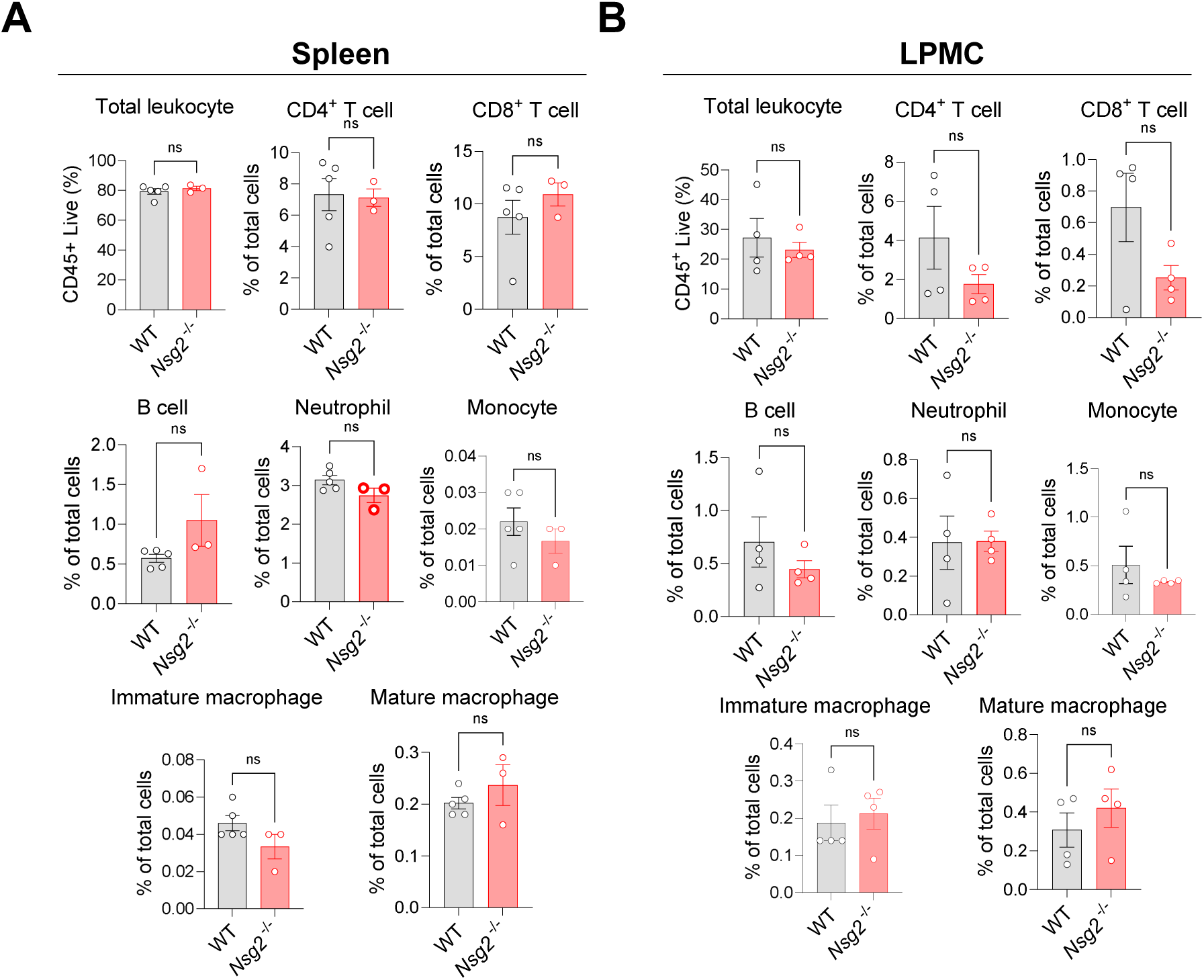
Baseline immune cell profiling in the spleen and colonic lamina propria of WT and *Nsg2*^-/-^ mice. Flow cytometric quantification of major immune cell populations in the (A) spleen and (B) colonic lamina propria (LPMC) of WT and *Nsg2*^-/-^ mice under steady-state conditions. Data are presented as a percentage of total live cells for each population. Each dot represents an individual mouse (n = 4 per group). Bars indicate group means ± SEM. Statistical significance was determined by unpaired two-tailed Student’s t-test. ns, not significant.

**Supplementary Figure S3.**
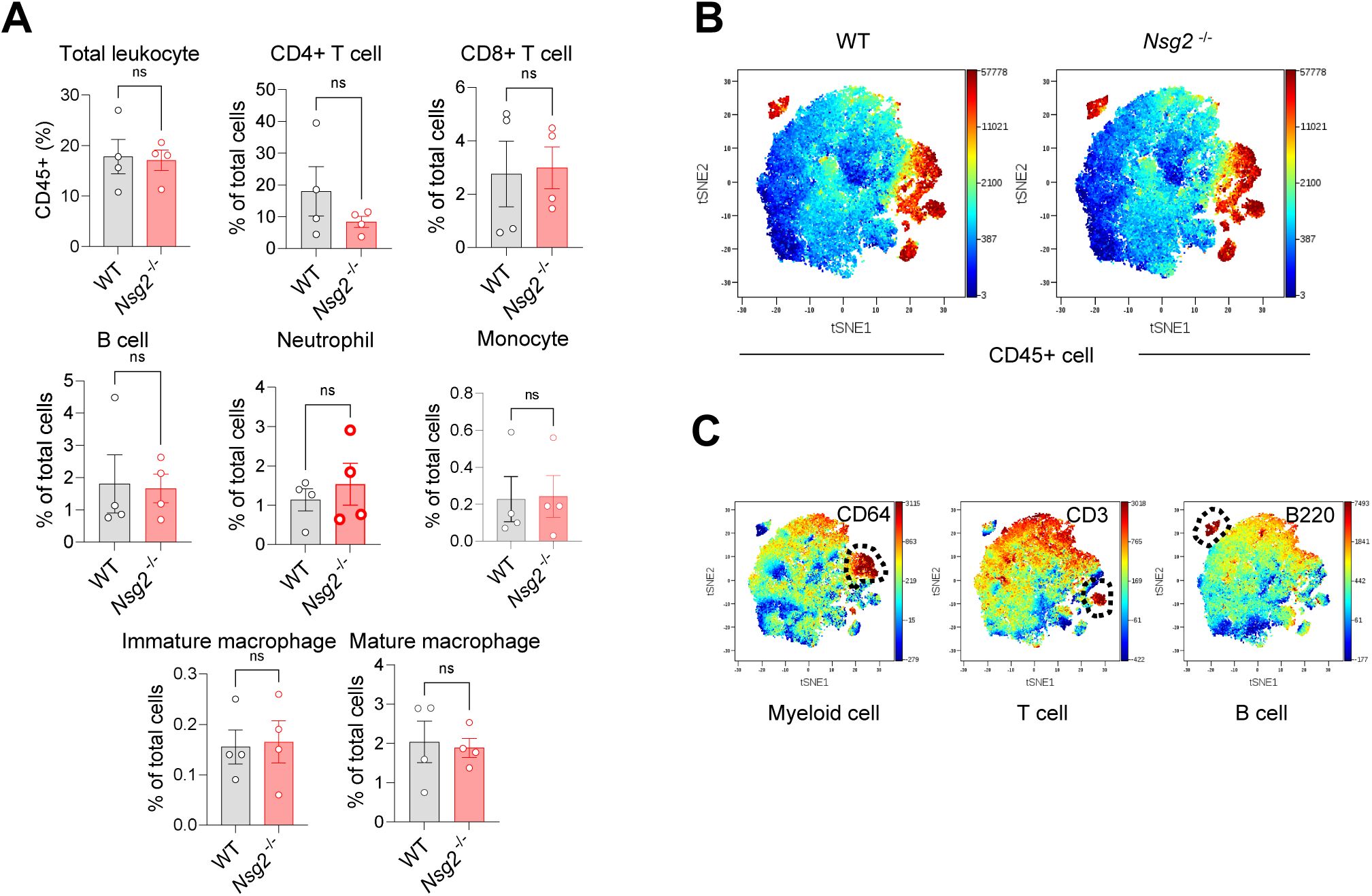
Comprehensive immune cell profiling in LPMC following AOM/DSS-induced CAC. (A) Flow cytometric quantification of major immune cell populations in colonic lamina propria from WT and *Nsg2*^-/-^ mice following AOM/DSS treatment. Data are presented as a percentage of total live cells for each population. Each dot represents an individual mouse (n = 4 per group). Horizontal lines indicate group means ± SEM. Statistical significance was determined by unpaired two-tailed Student’s t-test. \**p* < 0.05; ns, not significant. (B) Representative t-distributed Stochastic Neighbor Embedding (t-SNE) plots showing the overall distribution and clustering patterns of CD45^+^ immune cells from WT and *Nsg2*^-/-^ mice. Each plot represents pooled data from all mice within each genotype group. Colors indicate cell density, with red representing high-density regions and blue representing low-density regions. (C) High-resolution t-SNE visualization of distinct immune cell clusters identified through unsupervised machine learning analysis. Major cell populations are color-coded and annotated based on their characteristic surface marker expression profiles: myeloid cells (left), T cells (middle), and B cells (right). Black dots highlight specific cell clusters that show differential abundance between genotypes. Flow cytometry analysis was performed using standard protocols as described in the Methods section. t-SNE analysis was conducted using FlowJo software (Tree Star Inc., Ashland, OR, USA) with perplexity = 30 and 1000 iterations to ensure optimal clustering resolution.

**Supplementary Figure S4.**
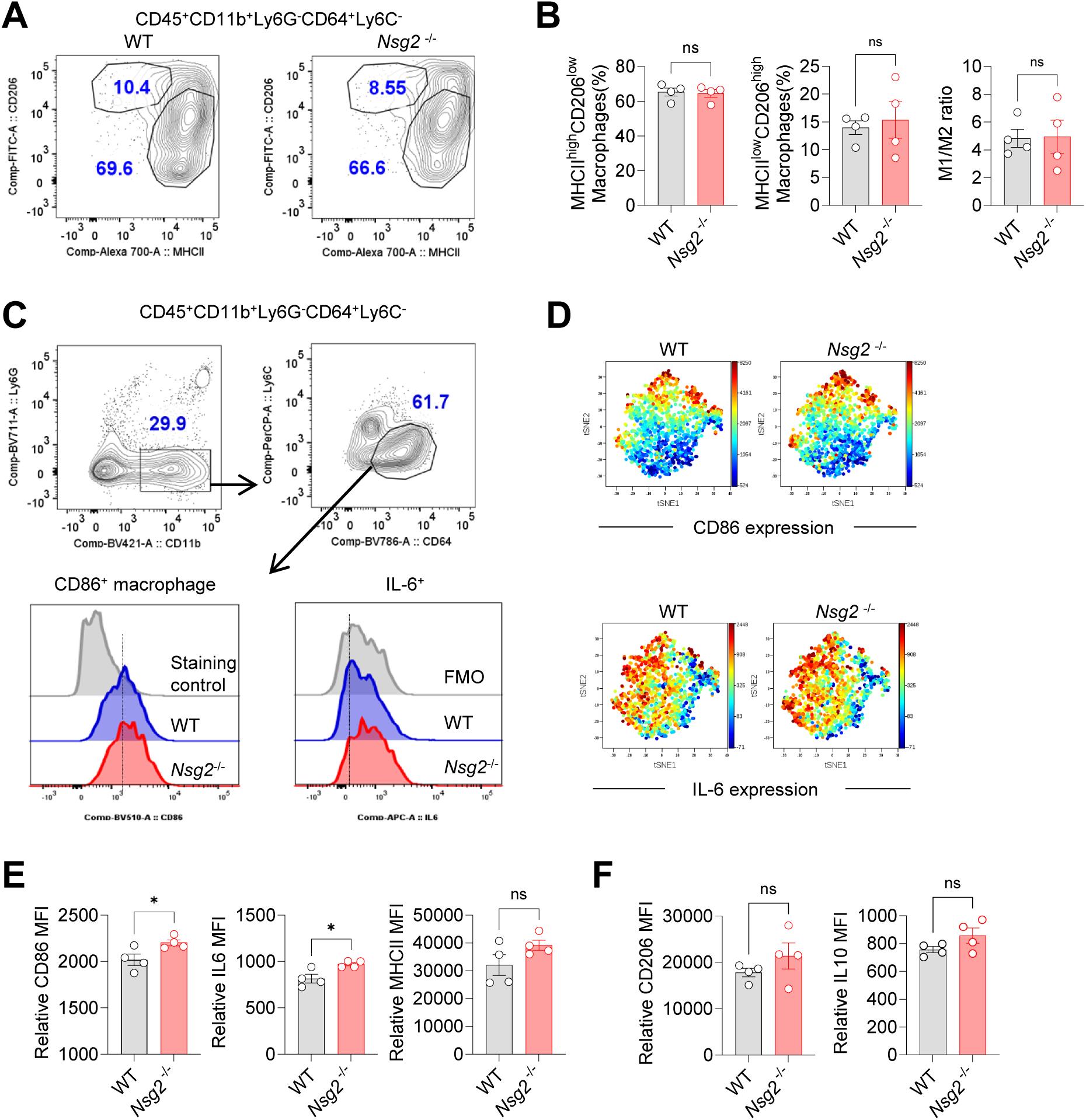
Detailed flow cytometric analysis of macrophage subsets and functional markers in AOM/DSS-induced CAC. (A) Representative frequency plots of M1 (MHCII^high^CD206^low^) and M2 (MHCII^high^CD206^high^) macrophages gated on CD45⁺CD11b⁺Ly6G⁻CD64⁺Ly6C^-^ cells. (B) Percentages of M1 (left) and M2 (middle) macrophage in LPMC, along with the calculated M1/M2 ratio (right). (C) Representative flow cytometry plots showing the gating strategy for colonic macrophages (CD45⁺CD11b⁺Ly6G⁻CD64⁺Ly6C^-^ cells), with quantification of CD86^+^ and IL-6^+^ subsets. The upper panel illustrates sequential gating of colonic macrophages from total LPMCs. The lower panel shows overlaid histograms comparing CD86 (left) and IL-6 (right) expression in WT and *Nsg2*^-/-^ mice. Staining controls (isotype and fluorescence minus one, FMO) are shown in gray. (D) tSNE plots visualizing single-cell expression of CD86 and IL-6 in colonic macrophages from WT and *Nsg2*^-/-^ mice. (E) Relative mean fluorescence intensity (MFI) of CD86, IL-6, and MHCII in M1 macrophages. (F) Relative MFI of CD206 and IL-10 expression in M2 macrophages. Data are presented as mean ± SEM. \**p* < 0.05; ns, not significant (unpaired two-tailed Student’s t-test).

## References

1. Lee, J., Im, J.P., Han, K., Kim, J., Lee, H.J., Chun, J., and Kim, J.S. (2020). Changes in Direct Healthcare Costs before and after the Diagnosis of Inflammatory Bowel Disease: A Nationwide Population-Based Study. Gut Liver 14, 89–99. 10.5009/gnl19023.

2. Im, J.P., Ye, B.D., Kim, Y.S., and Kim, J.S. (2018). Changing treatment paradigms for the management of inflammatory bowel disease. Korean J Intern Med 33, 28–35. 10.3904/kjim.2017.400.

3. Lee, H.S., Lobbestael, E., Vermeire, S., Sabino, J., and Cleynen, I. (2021). Inflammatory bowel disease and Parkinson’s disease: common pathophysiological links. Gut 70, 408–417. 10.1136/gutjnl-2020-322429.

4. Zhang, B., Wang, H.H.E., Bai, Y.M., Tsai, S.J., Su, T.P., Chen, T.J., Wang, Y.P., and Chen, M.H. (2021). Inflammatory bowel disease is associated with higher dementia risk: a nationwide longitudinal study. Gut 70, 85–91. 10.1136/gutjnl-2020-320789.

5. Günther, C., Rothhammer, V., Karow, M., Neurath, M., and Winner, B. (2021). The Gut-Brain Axis in Inflammatory Bowel Disease-Current and Future Perspectives. Int J Mol Sci 22. ARTN 887010.3390/ijms22168870.

6. Fung, T.C., Olson, C.A., and Hsiao, E.Y. (2017). Interactions between the microbiota, immune and nervous systems in health and disease. Nat Neurosci 20, 145–155. 10.1038/nn.4476.

7. Bonaz, B., Sinniger, V., and Pellissier, S. (2021). Therapeutic Potential of Vagus Nerve Stimulation for Inflammatory Bowel Diseases. Front Neurosci-Switz 15. ARTN 65097110.3389/fnins.2021.650971.

8. Chander, P., Kennedy, M.J., Winckler, B., and Weick, J.P. (2019). Neuron-Specific Gene 2 (NSG2) Encodes an AMPA Receptor Interacting Protein That Modulates Excitatory Neurotransmission. Eneuro 6. Artn 0292-18.201810.1523/Eneuro.0292-18.2018.

9. Nyuyki, K.D., and Pittman, Q.J. (2015). Toward a better understanding of the central consequences of intestinal inflammation. Ann Ny Acad Sci 1351, 149–154. 10.1111/nyas.12935.

10. Koh, S.J., Kim, J.M., Kim, I.K., Ko, S.H., and Kim, J.S. (2014). Anti-inflammatory mechanism of metformin and its effects in intestinal inflammation and colitis-associated colon cancer. J Gastroen Hepatol 29, 502–510. 10.1111/jgh.12435.

11. Koh, S.J., Choi, Y., Kim, B.G., Lee, K.L., Kim, D.W., Kim, J.H., Kim, J.W., and Kim, J.S. (2016). Matricellular Protein Periostin Mediates Intestinal Inflammation through the Activation of Nuclear Factor κB Signaling. Plos One 11. ARTN e0149652 10.1371/journal.pone.0149652.

12. Bain, C.C., and Mowat, A.M. (2014). Macrophages in intestinal homeostasis and inflammation. Immunol Rev 260, 102–117. 10.1111/imr.12192.

13. Hegarty, L.M., Jones, G.R., and Bain, C.C. (2023). Macrophages in intestinal homeostasis and inflammatory bowel disease. Nat Rev Gastro Hepat 20, 538–553. 10.1038/s41575-023-00769-0.

14. Yoo, B.B., and Mazmanian, S.K. (2017). The Enteric Network: Interactions between the Immune and Nervous Systems of the Gut. Immunity 46, 910–926. 10.1016/j.immuni.2017.05.011.

15. Wang, H.Z., Foong, J.P.P., Harris, N.L., and Bornstein, J.C. (2022). Enteric neuroimmune interactions coordinate intestinal responses in health and disease. Mucosal Immunol 15, 27–39. 10.1038/s41385-021-00443-1.

16. Hanahan, D., and Monje, M. (2023). Cancer hallmarks intersect with neuroscience in the tumor microenvironment. Cancer Cell 41, 573–580. 10.1016/j.ccell.2023.02.012.

17. Battaglin, F., Jayachandran, P., Strelez, C., Lenz, A., Algaze, S., Soni, S., Lo, J.H., Yang, Y., Millstein, J., Zhang, W., et al. (2022). Neurotransmitter signaling: a new frontier in colorectal cancer biology and treatment. Oncogene 41, 4769–4778. 10.1038/s41388-022-02479-4.

18. Venkatesh, H., and Monje, M. (2017). Neuronal Activity in Ontogeny and Oncology. Trends Cancer 3, 89–112. 10.1016/j.trecan.2016.12.008.

19. McCafferty, J., Mühlbauer, M., Gharaibeh, R.Z., Arthur, J.C., Perez-Chanona, E., Sha, W., Jobin, C., and Fodor, A.A. (2013). Stochastic changes over time and not founder effects drive cage effects in microbial community assembly in a mouse model. Isme J 7, 2116–2125. 10.1038/ismej.2013.106.

20. Ridaura, V.K., Faith, J.J., Rey, F.E., Cheng, J.Y., Duncan, A.E., Kau, A.L., Griffin, N.W., Lombard, V., Henrissat, B., Bain, J.R., et al. (2013). Gut Microbiota from Twins Discordant for Obesity Modulate Metabolism in Mice. Science 341, 1079–U1049. 10.1126/science.1241214.

21. Geirnaert, A., Steyaert, A., Eeckhaut, V., Debruyne, B., Arends, J.B.A., Van Immerseel, F., Boon, N., and Van de Wiele, T. (2014). , a butyrate producer with probiotic potential, is intrinsically tolerant to stomach and small intestine conditions. Anaerobe 30, 70–74. 10.1016/j.anaerobe.2014.08.010.

22. Boesmans, L., Valles-Colomer, M., Wang, J., Eeckhaut, V., Falony, G., Ducatelle, R., Van Immerseel, F., Raes, J., and Verbekem, K. (2018). Butyrate Producers as Potential Next-Generation Probiotics: Safety Assessment of the Administration of to Healthy Volunteers. Msystems 3. ARTN e00094-1810.1128/mSystems.00094-18.

23. Vacca, M., Celano, G., Calabrese, F.M., Portincasa, P., Gobbetti, M., and De Angelis, M. (2020). The Controversial Role of Human Gut Lachnospiraceae. Microorganisms 8. ARTN 57310.3390/microorganisms8040573.

24. Martin-Gallausiaux, C., Marinelli, L., Blottière, H.M., Larraufie, P., and Lapaque, N. (2021). SCFA: mechanisms and functional importance in the gut. P Nutr Soc 80, 37– 49. 10.1017/S0029665120006916.

25. Aggeletopoulou, I., and Triantos, C. (2024). Microbiome Shifts and Their Impact on Gut Physiology in Irritable Bowel Syndrome. Int J Mol Sci 25. ARTN 1239510.3390/ijms252212395.

26. Wei, L., Singh, R., Ro, S., and Ghoshal, U.C. (2021). Gut microbiota dysbiosis in functional gastrointestinal disorders: Underpinning the symptoms and pathophysiology. Jgh Open 5, 976–987. 10.1002/jgh3.12528.

27. Mayer, E.A., Savidge, T., and Shulman, R.J. (2014). Brain-Gut Microbiome Interactions and Functional Bowel Disorders. Gastroenterology 146, 1500–1512. 10.1053/j.gastro.2014.02.037.

28. Ghoshal, U., Shukla, R., Srivastava, D., and Ghoshal, U.C. (2016). Irritable Bowel Syndrome, Particularly the Constipation-Predominant Form, Involves an Increase in, Which Is Associated with Higher Methane Production. Gut Liver 10, 932–938. 10.5009/gnl15588.

29. Gandhi, A., Shah, A., Jones, M.P., Koloski, N., Talley, N.J., Morrison, M., and Holtmann, G. (2021). Methane positive small intestinal bacterial overgrowth in inflammatory bowel disease and irritable bowel syndrome: A systematic review and meta-analysis. Gut Microbes 13. Artn 193331310.1080/19490976.2021.1933313.

30. Nogal, A., Valdes, A.M., and Menni, C. (2021). The role of short-chain fatty acids in the interplay between gut microbiota and diet in cardio-metabolic health. Gut Microbes 13. 10.1080/19490976.2021.1897212.

31. Yurimoto, H., Kato, N., and Sakai, Y. (2005). Assimilation, dissimilation, and detoxification of formaldehyde, a central metabolic intermediate of methylotrophic metabolism. Chem Rec 5, 367–375. 10.1002/tcr.20056.

32. Chen, N.H., Djoko, K.Y., Veyrier, F.J., and McEwan, A.G. (2016). Formaldehyde Stress Responses in Bacterial Pathogens. Front Microbiol 7. ARTN 25710.3389/fmicb.2016.00257.

33. Vorholt, J.A., Chistoserdova, L., Stolyar, S.M., Thauer, R.K., and Lidstrom, M.E. (1999). Distribution of tetrahydromethanopterin-dependent enzymes in methylotrophic bacteria and phylogeny of methenyl tetrahydromethanopterin cyclohydrolases. J Bacteriol 181, 5750–5757. Doi 10.1128/Jb.181.18.5750-5757.1999.

34. Warren, M.J., Raux, E., Schubert, H.L., and Escalante-Semerena, J.C. (2002). The biosynthesis of adenosylcobalamin (vitamin B). Nat Prod Rep 19, 390–412. 10.1039/B108967f.

35. Roth, J.R., Lawrence, J.G., and Bobik, T.A. (1996). Cobalamin (coenzyme B-12): Synthesis and biological significance. Annu Rev Microbiol 50, 137–181. DOI 10.1146/annurev.micro.50.1.137.

